# Modeling the Diverse Effects of Divisive Normalization on Noise Correlations

**DOI:** 10.1101/2022.06.08.495145

**Authors:** Oren Weiss, Hayley A. Bounds, Hillel Adesnik, Ruben Coen-Cagli

**Affiliations:** Department of Systems and Computational Biology, Albert Einstein College of Medicine; Helen Wills Neuroscience Institute, University of California, Berkeley; Department of Molecular and Cell Biology, University of California, Berkeley; Dominick P. Purpura Department of Neuroscience, Albert Einstein College of Medicine; Department of Ophthalmology and Visual Sciences, Albert Einstein College of Medicine

## Abstract

Divisive normalization, a prominent descriptive model of neural activity, is employed by theories of neural coding across many different brain areas. Yet, the relationship between normalization and the statistics of neural responses beyond single neurons remains largely unexplored. Here we focus on noise correlations, a widely studied pairwise statistic, because its stimulus and state dependence plays a central role in neural coding. Existing models of covariability typically ignore normalization despite empirical evidence suggesting it affects correlation structure in neural populations. We therefore propose a pairwise stochastic divisive normalization model that accounts for the effects of normalization and other factors on covariability. We first show that normalization modulates noise correlations in qualitatively different ways depending on whether normalization is shared between neurons, and we discuss how to infer when normalization signals are shared. We then apply our model to calcium imaging data from mouse primary visual cortex (V1), and find that it accurately fits the data, often outperforming a popular alternative model of correlations. Our analysis indicates that normalization signals are often shared between V1 neurons in this dataset. Our model will enable quantifying the relation between normalization and covariability in a broad range of neural systems, which could provide new constraints on circuit mechanisms of normalization and their role in information transmission and representation.

**Author Summary:** Cortical responses are often variable across identical experimental conditions, and this variability is shared between neurons (noise correlations). These noise correlations have been studied extensively to understand how they impact neural coding and what mechanisms determine their properties. Here we show how correlations relate to divisive normalization, a mathematical operation widely adopted to describe how the activity of a neuron is modulated by other neurons via divisive gain control. We introduce the first statistical model of this relation. We extensively validate the model and investigate parameter inference in synthetic data. We find that our model, when applied to data from mouse visual cortex, outperforms a popular model of noise correlations that does not include normalization, and it reveals diverse influences of normalization on correlations. Our work demonstrates a framework to measure the relation between noise correlations and the parameters of the normalization model, which could become an indispensable tool for quantitative investigations of noise correlations in the wide range of neural systems that exhibit normalization.

## 1 Introduction

Neurons in the sensory cortices of the brain exhibit substantial response variability across identical experimental trials [1, 2]. These fluctuations in activity are often shared between pairs of simultaneously recorded neurons, called noise correlations [3]. Because the presence of these correlations can constrain the amount of information encoded by neural populations and impact behavior [4–13], noise correlations have been widely studied. This work has also revealed that correlations are often modulated by stimulus and state variables [3, 14], and therefore can play an important role in computational theories of sensory coding. For instance, noise correlations could emerge from neurons performing Bayesian inference and reflect the statistics of sensory inputs [15–17, 132] and prior expectations [18–20]. From a mechanistic point of view, such a statistical structure of noise correlations poses strong constraints on circuit models of cortical activity [21–25]. To better understand the functional impact and underlying mechanisms of noise correlations on neural coding and behavior, we need to be able to quantitatively characterize and interpret how noise correlations in neural populations are affected by experimental variables.

For this reason, successful descriptive models of neural activity have been developed to capture noise correlations [26–33]. However, none of those models considers divisive normalization [34–36], an operation observed in a wide range of neural systems [37–39] which has also been implicated in modulating the structure of noise correlations. Experimental phenomena that are accompanied by changes in noise correlations, including contrast saturation [40], surround suppression [41–43], and attentional modulations of neural activity [44–46] have been successfully modeled using divisive normalization [35, 47–49], although those models only captured average firing rates of individual neurons. Additionally, some numerical simulation studies have shown how normalization can affect noise correlations [50, 51]. These results indicate that it is important to quantify the relative contribution of normalization and other factors to modulation of noise correlations in experimental data.

We propose a stochastic normalization model, the pairwise Ratio of Gaussians (RoG), to capture the across-trial joint response statistics for pairs of simultaneously recorded neurons. This builds on our previous method that considered the relationship between normalization and single-neuron response variability (hence we refer to it as the independent RoG; [52]). In these RoG models, neural responses are described as the ratio of two random variables: the numerator, which represents excitatory input to the neuron, and the denominator (termed normalization signal), which represents the suppressive effect of summed input from a pool of neurons [34, 53]. Our pairwise RoG allows for the numerators and denominators that describe the individual responses to be correlated across pairs; in turn, these correlations induce correlations between the ratio variables (i.e., the model neurons’ activity; Figure 1a). In this paper, we derive and validate a bivariate Gaussian approximation to the joint distribution of pairwise responses, which greatly simplifies the problem of fitting the model and interpreting its behavior. The model provides a mathematical relationship between noise correlations and normalization, which predicts qualitatively different effects of normalization on noise correlations, depending on the relative strength and sign of the correlation between numerators and between denominators. This could explain the diversity of modulations of noise correlations observed in past work [45, 51, 54]. To provide practical guidance for data-analytic applications of our model, we investigate the accuracy and stability of parameter inference, and illustrate the conditions under which our pairwise RoG affords better estimates of single-trial normalization signals compared to the independent RoG. We then demonstrate that the model fits accurately pairwise responses recorded in the mouse primary visual cortex (V1), and often outperforms a popular alternative that ignores normalization. In our dataset, we find that when the correlation parameter between denominators is significantly different from zero, it is positive, indicating that those pairs share their normalization signals.

Our results highlight the importance of modeling the relation between normalization and covariability to interpret the rich phenomenology of noise correlations. Our model and code provide a data-analytic tool that will allow researchers to further investigate such a relationship, and to quantitatively evaluate predictions made by normative and mechanistic models regarding the role of correlated variability and normalization in neural coding and behavior.

## 2 Methods

### 2.1 Generative Model – Pairwise Ratio of Gaussians (RoG)

Here we describe in detail the RoG model and derive the Gaussian approximation to the ratio variable. Note that our RoG is entirely different from those used in [55] and in [47], despite the same acronym: [55] refers to the distribution obtained from the ratio between two Gaussian distributions, whereas we refer to the random variable that results from the ratio of two Gaussian variables. [47] does not refer to probability distributions at all, but rather to a surround suppression model in which the center and surround mechanisms are characterized by Gaussian integration over their respective receptive fields, while the RoG considered here is a model of neural covariability in general.

We build from the standard normalization model [34, 53] which computes the across-trial average neural response (e.g., firing rate) as the ratio between a driving input to a neuron (*N*) and the summed input from a pool of nearby neurons (*D*):

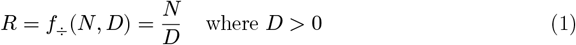

Where *f*_÷_ is the division operator; this functional notation is convenient for later derivations in which we consider the derivative of division. Our goal is to model the joint activity of pairs of neurons, so we extend the normalization model by considering two model neurons *R*_1_, *R*_2_. Since we are interested in trial-to-trial variability, we assume that a pair of neural responses ***R***_*t*_ = (*R*_1_, *R*_2_)_*t*_ on a single trial *t* can be written as the element-wise ratio of two Gaussian random vectors, ***N***_*t*_ = (*N*_1_, *N*_2_)_*t*_ and ***D***_*t*_ = (*D*_1_, *D*_2_)_*t*_, with additive Gaussian random noise *η*_*t*_ = (*η*_1_, *η*_2_)_*t*_ to capture the residual (i.e., stimulus-independent) variability.

As detailed further below, the numerators of two neurons can be correlated, and similarly for the denominators. In general, there can be correlations between the numerators and denominators (e.g., (*N*_1_, *D*_2_) may be correlated), requiring us to consider the joint, four-dimensional Gaussian distribution for the vector (***N***_*t*_, ***D***_*t*_). However, in this paper we consider the simpler model in which ***N***_*t*_ and ***D***_*t*_ are independent and are each distributed according to their respective two-dimensional Gaussian distributions. This assumption allows for simplified mathematical derivations and is supported by our previous work which found that including a parameter for the correlation between *N* and *D* caused over-fitting to single-neuron data [52]. However, we have also derived the equations for the case that numerators and denominators are correlated (see subsection S3), and implemented them in the associated code toolbox, so that interested researchers can test if their data warrant the inclusion of those additional free parameters.

We therefore write the generative model for the pairwise RoG as:

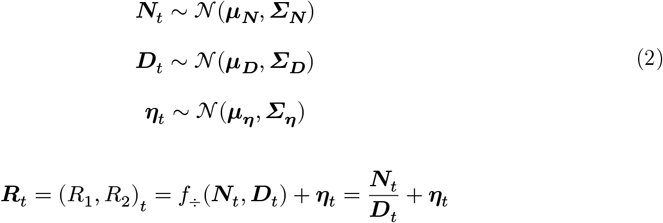

Where *f*_÷_ is applied element-wise, ***μ***_***N***_, ***μ***_***D***_ are the two-dimensional vectors of means of the numerator and denominator, respectively, ***Σ***_***N***_, ***Σ***_***D***_ are the respective 2 × 2 covariance matrices, and (***μ***_*η*_, ***Σ***_*η*_) is the mean and covariance matrix for the residual component of the model.

For the independent RoG, the ratio variable in general follows a Cauchy distribution whose moments are not well defined, but we used the result that when the denominator has negligible probability mass at values less than or equal to zero, the ratio distribution can be approximated by a Gaussian distribution with mean and variance that can be derived from a Taylor expansion [56–59]. This assumption is justified since the denominator is the sum of the non-negative responses from a pool of neurons [35] and is therefore unlikely to attain values less than or equal to zero.

For the pairwise extension, we can use the multivariate delta method (an application of a Taylor expansion) to compute the mean and covariance for the joint distribution of ratio variables [60] under the assumption that ***μ***_***D***_ > 0. We note that the true distribution of the ratio of bivariate or multivariate Gaussians vectors is unknown (although there is some work on ratios of complex Gaussian variables [135, 136]) and has higher-order statistics (e.g., skewness, kurtosis) that are not well approximated by an equivalent Gaussian. In this paper, we are interested in modeling the noise covariance as this is the most widely studied statistic in the field, and we show that the approximations we derive are very accurate (see Figure 1).

Future work could extend the model to account for these statistics by using higher-order terms in the Taylor expansion or a non-Gaussian copula.

To derive equations for the mean and covariance of the pairwise RoG, we use a Taylor expansion around the point (***μ***_***N***_, ***μ***_***D***_):

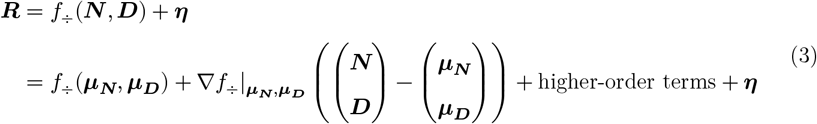

Using only the first order terms, we derive expressions for the mean and covariance matrix of the RoG:

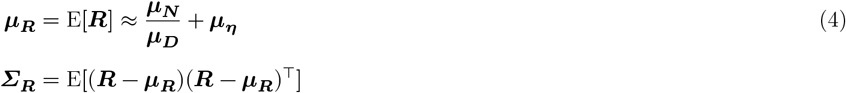

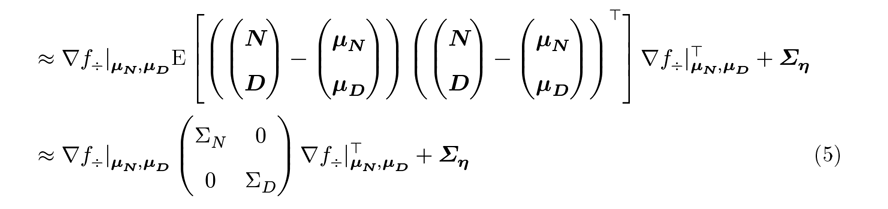

Note that the variance of the denominator influences the mean of the ratio variable through a second order term, hence it does not appear in Equation 4 (see [57] for the second order Taylor expansion for the mean of a ratio variable). From Equation 5, we can obtain expressions for the variance of each neuron in the pair and their covariance and correlation. First, we adopt the following notation to simplify the equations: let 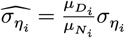 and let 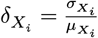.

Then:

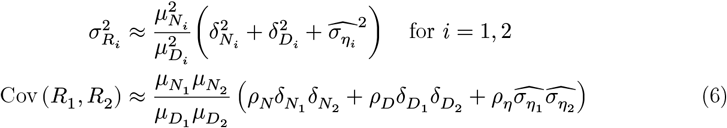

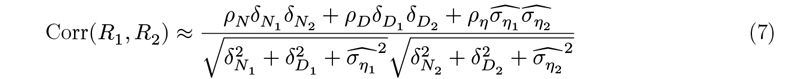

Equation 7 is commonly referred to as the formula for “spurious” correlation of ratios found when comparing ratios of experimental variables [61], and we further generalize this in subsection S3. To the extent that tuning similarity between neurons reflects similarity in the driving inputs, and that those driving inputs are variable, neurons with more similar tuning would have larger *ρ*_*N*_, which in turn implies larger noise correlations according to Equation 7. This is consistent with the widespread empirical observation that signal correlations and noise correlations are correlated [3].

### 2.2 Parametrization of the Pairwise RoG for Contrast Responses

In the form described so far, the pairwise RoG has 10 stimulus dependent parameters 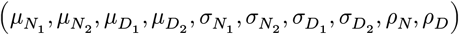 and 5 stimulus independent parameters 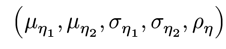 for the additive noise. For any stimulus condition, there are only five relevant measurements that can be derived from the neural data (the response means and variances for each neuron in a pair, and their correlation), so the model is over-parametrized. Therefore, to apply the RoG to neural data, we need to reduce the total number of parameters.

The generality of this model provides a procedure for converting a standard normalization model (i.e., a model for the mean response) into a RoG model that specifies both mean and (co)-variance. In this paper, we use the example of contrast-gain control, which has been widely used to model the response of single neurons and neural populations to visual input with varying contrast [35, 62–64]. By adapting such a model, we can reduce the stimulus dependence of the means of the numerator and denominator 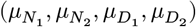. In the contrast-gain control model, the neural response as a function of contrast *c* (0 − 100%) is computed as a “hyperbolic ratio” [35, 62]:

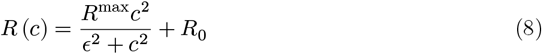

Where *R*^max^ is the maximum response rate, ϵ is the semi-saturation constant (the contrast at which *R* (ϵ) = *R*^max^/2) to prevent division by 0, and *R*_0_ is the spontaneous activity of the neuron (the response at 0% contrast). We can convert this standard model into an RoG by setting the mean of the numerator and denominator in the RoG to the numerator and denominator in this equation:

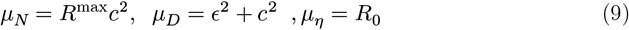

By using this functional form, we can substitute the stimulus dependent parameters of the 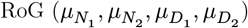 with the stimulus independent parameters 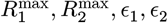.Another model simplification is to assume that individual neural variability and mean neural response are related by a power function as has been observed in the visual cortex [65– 67]:

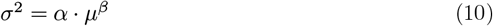

This parametrization allows the Fano Factor (the ratio of the variance to the mean) to vary with stimulus input (as long as *β* ≠ 1) and for both over-dispersion (Fano factor > 1) and under-dispersion (Fano factor < 1). Moreover, as with the mean, the four stimulus dependent variance parameters of the model 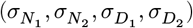 can be replaced with four pairs of stimulus independent parameters **(α**_***N***_, **β**_***N***_, **α**_***D***_, **β**_***D***_). Lastly, in principle, the parameters controlling correlation (*ρ*_*N*_, *ρ*_*D*_) can vary with stimulus conditions but for computational simplicity we assume that (*ρ*_*N*_, *ρ*_*D*_) are stimulus independent. However, even with this assumption, our model can capture stimulus dependent noise correlations (Figure 5) as often observed *in vivo* [40, 68, 69].

### 2.3 Fitting the RoG to Data

We optimize the values of the parameters, given a dataset, by maximum likelihood estimation. In this paper, we validate various properties of the pairwise RoG using synthetic data produced from the generative model. We will demonstrate the applicability of this model to neural data analysis by fitting the pairwise RoG to calcium imaging data (see subsection 2.7).

Based on our previous discussion, we assume that the model parameters (collectively denoted Θ) are stimulus independent. We consider our dataset {***R***_*t*_(*s*)} where *s* is the stimulus and *t* indexes the trial. We assume that, for each stimulus, our data is independent and identically distributed according to 𝒩(**μ**_***R***_(*s*), ***Σ***_***R***_ (*s*); Θ), and that data is independent across stimuli. We can therefore compute the negative log-likelihood of the data using the following equation (see subsection S2 for derivation):

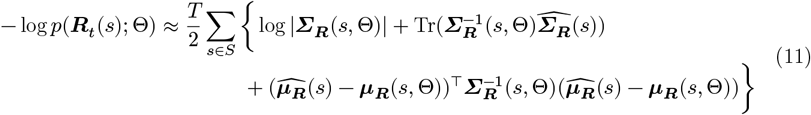

Where 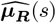 and 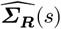 are the empirical mean and covariance across trials computed from the data.

In practice, we have found that it is computationally faster to first optimize the parameters for each neuron in the pair separately (which is equivalent to fitting the independent RoG model), and then optimize the correlation parameters (i.e., the *ρ* parameters) with the single-neuron model parameters fixed. This two-step optimization process is referred to as the inference functions for marginals method in the copula literature, and is known to be mathematically equivalent to maximum likelihood estimation for Gaussian copulas [70], which is the case we consider here. This points to an extension or alternative to the pairwise RoG that considers the bivariate distribution to be some non-Gaussian copula with Gaussian marginals, which we leave for future work. We assumed that the pairwise distribution is Gaussian for computational simplicity, but others have used non-Gaussian copulas to model neural populations [71].

#### 2.3.1 Cross-validated Goodness of Fit

To measure the quality of model fit, we used a cross-validated pseudo-*R*^2^ measure [72], as follows. During fitting, we divided the recording trials for each pair and for each stimulus into training and test sets (for simulation studies, we used two or ten-fold cross validation; for the calcium analysis we used leave-one-out cross-validation). We then fit the parameters of the model for each training set and used the following equation to assess the model prediction on the held-out data:

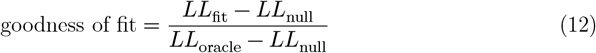

Where *LL*_fit_ is the negative log-likelihood (using Equation 11) for the test data using the optimized parameters, *LL*_null_ is the negative log-likelihood of the data assuming that there is no modulation of the responses by stimulus contrast, and *LL*_oracle_ is the negative log-likelihood of the data using the empirical mean, variance, and covariance of the training data per stimulus condition. The reported goodness of fit score is the median across all training and test splits of the computed score (Equation 12). Because of this cross-validation, goodness of fit values can be < 0 (the fit model is worse than the null model) or > 1 (the fit model performance is better than the oracle).

#### 2.3.2 Quantifying the Accuracy of the Estimated Correlation Parameters

As we are interested in interpreting the correlation model parameters (*ρ*_*N*_, *ρ*_*D*_), we need to assess the accuracy of the maximum likelihood estimator. For simulations, we directly compare the estimated *ρ* values to the true values used to generate the data. For real neural data, however, we do not have access to the true values: instead, we compute confidence intervals. To do so, we perform a bootstrap fit procedure: given a set of pairwise neural responses {***R***_*t*_(*s*)} with *T* simultaneously recorded trials, we sample these trials with replacement *T* times and then fit the pairwise RoG using the resampled set of neural responses as our observations. Repeating this procedure for a large number of samples (in the analysis in Sections 3.2 and 3.4 we used 1000 bootstrap samples) gives us sets of fit *ρ*_*N*_, *ρ*_*D*_, which we use to compute a 90% confidence interval. Using the synthetic data, we validate these confidence intervals by measuring the empirical coverage probability and comparing to the nominal confidence level. These confidence intervals allow one to quantify the accuracy of the *ρ* parameter estimates. We then demonstrate one possible use of the confidence intervals, with an application focused specifically on the sign of the *ρ* parameters.

### 2.4 Model Comparison

We compared the pairwise RoG to a modified version of the modulated Poisson model [66], using Gaussian noise instead of Poisson [73]. We call this model the Modulated Gaussian (MG). The original model is a compound Poisson-Gamma distribution, in which the Poisson rate parameter is the product of the mean tuning curve and a random gain variable that is Gamma distributed. The parameters of the Gamma distribution depend on the mean tuning curve (*f*(*s*)) and the variance of the gain variable (*σ*_*G*_). Additionally, there are two sources contributing to (tuned) covariability: the correlation between the multiplicative gains (*ρ*_*G*_), and the correlation between the Poisson processes (*ρ*_*P*_). For the modulated Gaussian model, we use a bivariate Gaussian distribution whose moments (i.e., mean, variance, and covariance) are parametrized according to the moments of the modulated Poisson model. We made this modification to the modulated Poisson model for two main reasons. First, because we are examining continuously valued fluorescence traces as opposed to discrete spike count data, a continuous distribution is more appropriate for analysis. Second, the original modulated Poisson, while including a parametrization of the noise covariance between neurons, has no simple closed form for the bivariate distribution, which complicates the comparison of goodness of fit between the two models. By using a bivariate Gaussian distribution, we can more directly compare this model to our proposed pairwise RoG.

More explicitly, the pairwise neural responses are distributed as:

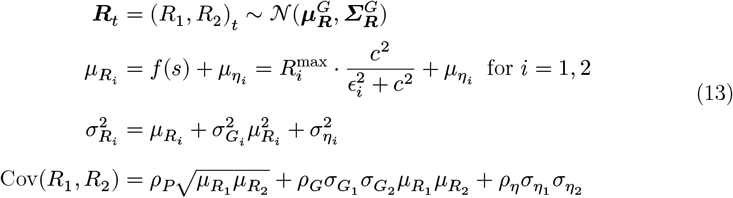

where we assume the mean tuning cure is the contrast-response curve (Equation 8), 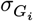 is the standard deviation of the multiplicative gain for neuron *i*, and *ρ*_*G*_ is the correlation between the multiplicative gains. *ρ*_*P*_ is no longer interpreted as the point process correlation; instead, *ρ*_*P*_ controls the portion of the tuned covariability that is independent of the shared gain. As with the RoG, we also model the untuned variability *η* as additive bivariate Gaussian noise. We then fit the model parameters to data by minimizing the negative log-likelihood (Equation 11, with 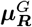 and 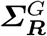 defined in Equation 13). As with the pairwise RoG, we use cross-validation to account for model complexity and compute the goodness of fit scores using Equation 12. An extension to this model was recently proposed that incorporates normalization by assuming the rate parameter is a ratio term in which the denominator is a Gaussian random variable, then deriving moments of the distribution for optimization [74].

However, this model does not currently account for noise correlations, so we chose to instead adapt the Poisson-Gamma model.

The most relevant difference between RoG and MG is in how each model accounts for the effect of normalization on (co)variability. In the RoG, normalization directly influences variability by division operating on random variables. This creates flexible dependencies between the mean firing rate, individual neuron variability and shared covariability. In the MG, normalization influences the gain of neurons through the interaction between the mean firing rate (i.e., the standard normalization model) and the gain parameter *σ*_*G*_, which is assumed be a slowly fluctuating source of variability that scales how the mean firing rate effects variability. In this way, the normalization signal for the MG is a deterministic factor. The MG is therefore a simpler model that can only account for overdispersion, whereas the RoG allows for both overdispersion and underdispersion, and for diverse patterns of covariability, as we show in Figure 5, albeit at the cost of additional parameters.

### 2.5 Inference of Single Trial Normalization from Measured Neural Activity

Because of the probabilistic formulation of the RoG, we can use Bayes theorem to compute the posterior probability of the normalization variable ***D***_*t*_ in a single trial, given the observed neural responses ***R***_*t*_:

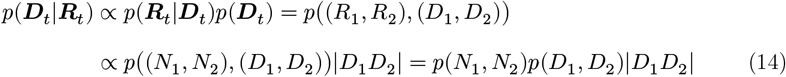

Where multiplication by |*D*_1_*D*_2_| occurs due to the change of variables formula for probability density functions. From this distribution, we can find the maximum a posteriori (MAP) estimates of the normalization strength in a single trial by differentiating the posterior distribution with respect the denominator variables and finding the maxima by setting the partial derivatives to 0. For ease of computation, we solve the equivalent problem of finding the zeros of the partial derivatives of the negative logarithm of the posterior distribution. In our previous work [52], we found that, when subtracting the mean additive noise from the simulated activity, the MAP estimate remained unbiased. Thus, for simplicity, we assume that we can subtract off the mean spontaneous activity and consider instead the posterior *p* (***D***|***R*** − *μ*_*η*_). To obtain an estimate for the denominator strength ***D***_*t*_ we look at the partial derivatives of the negative log posterior with respect to *D*_1_, *D*_2_ and solve to obtain the MAP estimate. This procedure leads to a two-dimensional system of bivariate quadratic equations:

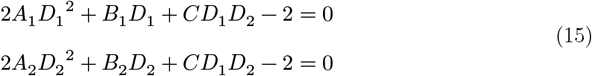

Where the coefficients *A*_1_, *A*_2_, *B*_1_, *B*_2_, *C* are functions of the parameters of the model (see subsection S4 for the derivation of Equation 15 and the full expressions of these coefficients).

A basic result from the algebra of polynomial systems (Bézout’s theorem) tells us that this system has four pairs of possibly complex valued solutions [75]. In fact, as solving this system amounts to solving a quartic equation in one variable, there exists an algebraic formula (with radicals) for solutions to this system as a function of the coefficients. This solution is too long to include here and uninformative but was found using the Symbolic Math toolbox from MATLAB and is included in our toolbox (subsection 2.8).

Because all the variables involved are real-valued, we are only interested in the existence of real solutions to this two-dimensional system. However, there is no theoretical guarantee that there will be any real solutions. In practice we take the real part of the algebraic solution to this system and find which pair of solutions 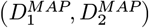 minimize the negative log posterior. Alternatively, we can consider finding the MAP by directly minimizing Equation 14 (see subsection S4) using numerical optimization. We have verified that, when real-valued solutions exist to Equation 15, these coincide with numerically minimizing Equation 14. However, as optimization of Equation 14 must be computed on a per-trial basis, it is far too time consuming to perform when there are many experimental trials, so we utilize the algebraic solution to Equation 15.

### 2.6 Generating Realistic Pairwise Neural Activity from the Model

To constrain our simulations to realistic parameter values for the contrast response function (Equation 8), we took the single-neuron best fit parameters to macaque V1 data analyzed in our previous work (for details see [52]) and created parameter pairs by considering all combinations (*N* = 11628 pairs) of these parameters. Using the generative model for the pairwise RoG (Equation 2) and the contrast-response parametrization (Equation 8), we can simulate single-trial neural activity from these parameter pairs and specific values for (*ρ*_*N*_, *ρ*_*D*_). These synthetic data allow us to explore properties of the pairwise model without having to exhaustively explore the full parameter space.

### 2.7 Data Collection and Processing

#### 2.7.1 Ethics Statement

All experiments on animals were conducted with approval of the Animal Care and Use Committee of the University of California, Berkeley.

#### 2.7.2 Animal Preparation

Data were collected from CaMKII-tTA;tetO-GCaMP6s mice [76], expressing GCaMP6s in cortical excitatory neurons. Mice were implanted with headplates and cranial windows over V1 [77]. Briefly, mice were anesthetized with 2% isoflurane and administered 2 mg/kg of dexamethasone and 0.5 mg/kg of buprenorphine. Animals were secured in a stereotaxic frame (Kopf) and warmed with a heating pad. The scalp was removed and the skull was lightly etched. A craniotomy was made over V1 using a 3.5 mm skin biopsy bunch. A cranial window, consisting of two 3 mm diameter circular coverslips glued to a 5 mm diameter circular coverslip, was placed onto the craniotomy, and secured into place with Metabond (C&B). Then a custom-made titanium headplate was secured via Metabond (C&B) and the animals were allowed to recover in a heated cage.

#### 2.7.3 Behavioral Task and Visual Stimuli

During imaging, mice were head-fixed in a tube, and were performing an operant visual detection task [78]. Briefly, mice were trained to withhold licking when no stimulus was present, and lick within a response window after stimulus presentation. Mice were water-restricted and given a water reward for correct detection. Visual stimuli were drifting sinusoidal gratings (2 Hz, 0.08 cycles/degree) presented for 500 ms followed by a 1000 ms response window. Stimuli were generated and presented using PsychoPy2 [79]. Visual stimuli were presented using a gamma corrected LCD monitor (Podofo, 25 cm, 1024x600 pixels, 60 Hz refresh rate) located 10 cm from the right eye. Contrast of gratings were varied between 7 different levels: {2, 8, 16, 32, 64, 80, 100}, except for 2 recording sessions in which contrast level 80% was omitted. This did not alter any of the analysis, allowing sessions to be combined into a single dataset.

#### 2.7.4 Calcium Imaging

Once they learned the task, mice started performing under the 2p microscope, and V1 was imaged via cranial window. Imaging was performed using a 2-photon microscope (Sutter MOM, Sutter Inc.), with a 20X magnification (1.0 NA) water-immersion objective (Olympus Corporation). Recordings were done in L2/3 in an 800 x 800 μm field of view, with 75-100 mW of 920 nm laser light (Chameleon; Coherent Inc). An electrically tunable lens (Optotune) was used to acquire 3 plane volumetric images at 6.36 Hz. Planes were 30 μm apart. Acquisition was controlled with ScanImage (Vidrio Technologies).

Calcium imaging data was motion-corrected and ROI extracted using suite2p [80], and all data was neuropil subtracted with a coefficient of 0.7 (we also analyzed data using neuorpil coefficients of 0.4 and 1, not shown, and see subsection S6 for additional analysis with deconvolved data; all the results presented in the main text were qualitatively similar across preprocessing methods).

#### 2.7.5 Data Processing

Processing of calcium imaging data was performed using custom MATLAB code. Fluorescence traces for individual trials and cells (average of the neuropil subtracted fluorescence across a ROI) consisted of 24 frames: 4 frames of pre-stimulus blank, followed by 3 frames of stimulus presentation and 17 frames of post-stimulus blanks corresponding to the response window for the behavioral task. In our analyses, we considered one extra frame to account for onset delays and calcium dynamics. Baseline fluorescence (*F*_0_) was computed as the median across pre-stimulus frames (1-5), and the stimulus evoked fluorescence (Δ*F*/*F*) was computed as the mean of the normalized fluorescence per frame ((*F* (*i*) − *F*_0_)/*F*_0_ for *F* (*i*) the fluorescence of frame *i*) across frames corresponding to stimulus response. Spontaneous Δ*F*/*F* was computed as above during blank trials. Cells were included in further analysis if the evoked response at the highest contrast was at least 2 standard deviations above the spontaneous mean fluorescence. Across 9 recording sessions, 295/8810 neurons met this inclusion criterion. This small percentage is due to sessions recorded using gratings with fixed spatial frequency and orientation; thus, included neurons are visually responsive and selective for this combination.

### 2.8 Code and Data Availability

Code for running model simulations and fitting model to experimental data was written in MATLAB and is publicly available on Github (https://github.com/oren-weiss/pairwiseRatioGaussians). Calcium imaging data is publicly available on Zenodo (https://zenodo.org/doi/10.5281/zenodo.8118187).

## 3 Results

We developed the pairwise Ratio of Gaussians model (RoG, Figure 1a) to quantify the relationship between normalization and response covariability (i.e., noise correlations) that has been suggested in empirical studies of neural activity in visual cortex [51, 54, 81, 82]. In the standard divisive normalization model (Equation 1), the mean response is computed as the ratio of the excitatory drive (numerator *N*) to a normalization signal summarizing inhibitory drive (denominator *D*). Our pairwise RoG considers a pair of neurons where each individual neural response is well characterized by the standard normalization equation with corresponding numerators (*N*_1_, *N*_2_) and denominators (*D*_1_, *D*_2_). We then assume that the numerators and denominators are bivariate Gaussian random vectors -which allows the possibility for correlations to exist among the numerators (denoted *ρ*_*N*_) and among the denominators (*ρ*_*D*_). From this, we derived equations for the mean responses and covariance matrix of the pair as a function of the numerators and denominators (Eqs. 4, 5). These equations depend on the Gaussian approximation to the ratio of two Gaussian random variables. We verified the validity of this approximation for the moments of interest (mean, variance, and covariance), by simulating the activity of pairs of neurons, and comparing the covariance (Figure 1b,c) and correlation (Figure 1d,e) of the true ratio distribution and of the approximate distribution (Eqs. 6, 7). The mean and variance are identical to the independent RoG model (a special case of the pairwise RoG, with numerators and denominators independent between neurons) which we validated previously [52].

**Figure 1:**
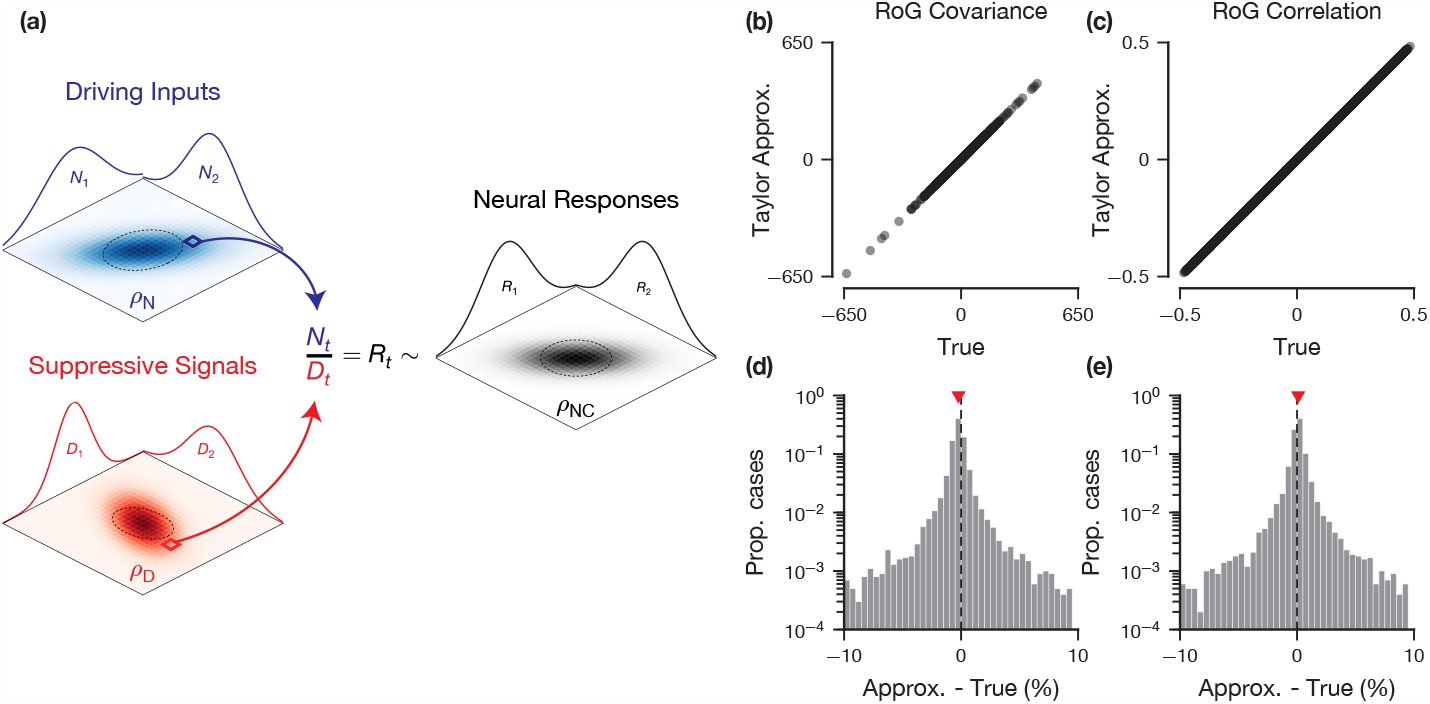
Definition and Validation of the Pairwise Ratio of Gaussians Model. (a) The pairwise RoG model describes pairs of neural responses (*R*_1_, *R*_2_), where each response is computed as the ratio of two stimulus-driven signals on a trial-by-trial basis: numerators (*N*_1_, *N*_2_), representing the driving inputs; and denominators (*D*_1_, *D*_2_), representing the suppressive signals. Across trials, the numerators and denominators are distributed according to bivariate Gaussian distributions with correlation coefficients (ρ_*N*_, ρ_*D*_), respectively. The resulting response distribution is approximately Gaussian with correlation coefficient ρ_*NC*_. (b-e) Comparison of the normal approximation we derived for the pairwise RoG noise covariance (b,c) and noise correlation (d,e) and the true values (estimated across 1*e*6 simulated trials) for 1*e*4 experiments (i.e., simulated pairs of neural responses). Each experiment used different model parameters and each trial was randomly drawn from the corresponding distribution. (b,d) scatter plot; (c,e) histogram of the percent difference between the Taylor approximation and the true value. The red marker indicates the median percent difference. Model parameters were drawn uniformly from the following intervals: μ_***N***_ ∈ [0, 100], μ_***D***_ ∈ [0.5, 1.5], ***μ***_*η*_ = 0, α_***N***_ = 1, α_***D***_ = 0.001, ***β***_***N***_, ***β***_***D***_ ∈ [1, 1.5], ***ρ***_*N*_, ***ρ***_*D*_ ∈ [−0.5, 0.5], 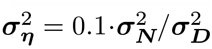, ***ρ***_*η*_ = 0. The ranges of the mean parameters were chosen to reproduce realistic firing rates of V1 cortical neurons, while the ***α, β*** parameters were chosen such that the variances of the *N* and *D* are relatively small and the probability that ***D*** ≤ 0 is negligible [56].

### 3.1 Modulations of Correlated Variability Depend on Sharing of Normalization

Within the RoG modeling framework, there are two main sources of response (co)-variability: the numerator (excitatory drive) and the denominator (normalization). Depending on the value of the corresponding *ρ* parameters, each of these sources of variability can be independent (ρ = 0) or shared (ρ ≠ 0), and therefore contribute differently to noise correlations. Consequently, understanding modulations of noise correlations in the pairwise RoG requires understanding how normalization and external stimuli affect the relative amplitude of these sources, and how the effects depend on whether those sources are shared.

First, we studied the relationship between normalization and noise correlations for the lowest contrast stimuli (Figure 2, yellow symbols). We define normalization strength for a given neuron as the mean of the denominator; for a fixed stimulus contrast, this is determined by the semi-saturation constant ϵ in Equation 8. When the normalization signals are positively correlated (shared normalization, *ρ*_*D*_ = 0.5) but the excitatory drive is independent (*ρ*_*N*_ = 0), increasing normalization strength tends to decrease the magnitude of noise correlations (Figure 2a). Conversely, when the normalization signals are independent (*ρ*_*D*_ = 0) but there is shared driving input (*ρ*_*N*_ = 0.5), the magnitude of noise correlations tends to increase with increasing normalization (Figure 2b). Intuitively, this is due to how the model partitions neuronal covariability into two sources and how normalization separately effects variability of these sources. As mentioned at the beginning of this section, these terms describe the correlations among the numerators and among the denominators. The correlations arising from the numerator are unaffected by normalization strength, while the correlations arising from the denominator tend to decrease with normalization strength. So, when *ρ*_*N*_ = 0, the noise correlations are solely due to the denominator cofluctuations and thus tend to decrease with normalization. However, when *ρ*_*D*_ = 0, the numerator covariability drives the response covariability, which is unchanged by normalization strength. The reason why we see increased noise correlations in this scenario is because normalization decreases individual neuronal response variability [52] so the proportional contribution of the numerator term increases. This is derived more completely later in this section. Therefore, increasing normalization strength reduces a source of noise correlations when normalization is shared, whereas it reduces a source of independent noise when normalization is not shared.

We found similar effects of normalization on noise correlations at high contrast levels (Figure 2, orange and red symbols) although the slope of the relationship became shallower as contrast increased. This is due to the saturating effect of the contrast response function (Equation 8): at high contrast, (co)fluctuations of the normalization signal across trials have relatively little effect on the responses of a neural pair, so the correlation in neural responses will be relatively unaffected by normalization strength. We also observed that the magnitude of noise correlations generally increased with contrast when the denominators were correlated and the numerators independent (Figure 2a and S3), whereas it decreased when the numerators were correlated (Figure 2b,c and S3). Importantly, for the analysis of normalization strength at fixed contrast, we used the contrast semi-saturation constants (Equation 8; i.e., a pure change in the denominator) as a measure of normalization strength. Conversely, increasing stimulus contrast increases both the numerator and denominator (Equation 8). This explains our observation that, even though normalization is stronger at higher contrast, noise correlations can be modulated in different ways by contrast and by normalization, because changing stimulus contrast also affects the numerator term.

Indeed, these results can be derived from examining the equation for the correlation in the pairwise RoG (Equation 7) rearranged as follows (see subsection S1 for more details):

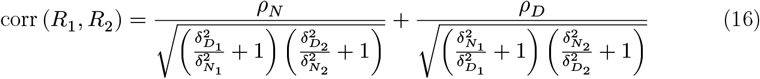

and by recognizing that the coefficient of variation of 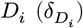 is a decreasing function of normalization strength (*μ*_*D*_ or *ϵ*), whereas the ratio 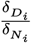 is *often* an increasing function of contrast.

We further analyze this equation, first in the case of changing normalization strength while keeping contrast fixed. The term proportional to *ρ*_*N*_ is an increasing function of the mean normalization strength since the denominator is a decreasing function of μ_*D*_. Conversely, the term proportional to *ρ*_*D*_ decreases with normalization since the denominator increases with μ_*D*_. In these two cases, the monotonic dependence of noise correlation on normalization strength is guaranteed by Equation 16 regardless of the specific parameter values. Similar patterns emerge when *ρ*_*N*_, *ρ*_*D*_ < 0, except the signs of the noise correlations are reversed (Figure S3). When the correlations of input and normalization signals are both different from zero, the relationship between noise correlation and normalization strength resembles a combination of the two previously described scenarios, and the specific parameter values determine which of the two terms in Equation 16 dominates. For instance, in our simulations with (*ρ*_*N*_ = 0.5, *ρ*_*D*_ = 0.5) (Figure 2c), the magnitude of noise correlations increased with normalization strength on average similar to (Figure 2b), indicating that the magnitude of the term proportional to *ρ*_*N*_ is usually larger than the magnitude of the *ρ*_*D*_ term, but this trend is not consistent, as evidenced by the increased spread of the scatter. When the input strength and normalization signal have opposite correlations (e.g., *ρ*_*N*_ = 0.5, *ρ*_*D*_ = −0.5), we obtained similar results; however, the magnitude of noise correlations was on average closer to 0 due to cancellation between the two sources of covariability (Figure S3).

A similar analysis of Equation 16 shows that the effects of stimulus contrast are opposite to those of a pure change of normalization strength, because 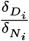 is often an increasing function of contrast but a decreasing function of normalization strength. Notice that we assumed there is no residual noise component (*η* ≡ 0), but all the analyses above remain valid when the amplitude of noise variance σ_*η*_ is relatively small compared to (δ_*N*_, *δ*_*D*_) (Equation S5).

**Figure 2:**
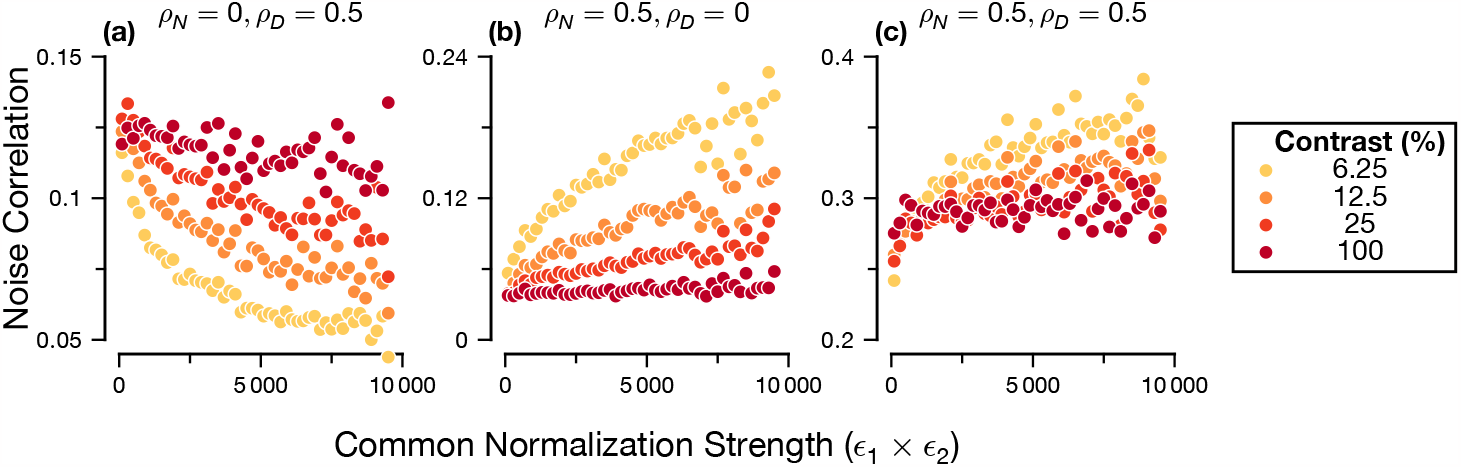
Relationship between Noise Correlations, Normalization Strength and Contrast Depends on the Source of Variability. Each panel shows, for a combination of *ρ*_*N*_, *ρ*_*D*_ specified in the panel title, the median noise correlation of all generated neural pairs binned according to ϵ_1_ϵ_2_, a contrast independent measure of the common normalization strength. Bins with less than 100 pairs were discarded. Neural responses were generated from the contrast-response parametrization (Equation 8). Noise correlation strength was computed across 1e3 simulated trials drawn from the pairwise RoG model. For each contrast level and combination of *ρ*_*N*_, *ρ*_*D*_, 1e5 simulated experiments were created. See Figure S4 for a more systematic exploration of the factors influencing modulation of noise correlations for two simulated pairs that matches the large scale experiment considered here. Model parameters were drawn uniformly from the following intervals: ***R***^max^ ∈ [5, 50], ϵ ∈ [10, 100], ***α***_***N***_, α_***D***_ ∈ [0.1, 1], ***β***_***N***_, ***β***_***D***_ ∈ [1, 2], ***η*** ≡ 0

In summary, our analysis shows that normalization and stimulus contrast can have diverse effects on noise correlations, depending on whether neurons share their normalization signals and on the interplay between multiple sources of variability.

### 3.2 Inference of Correlation Parameters

The above analysis demonstrates that the relationship between noise correlations and normalization depends on how this correlated variability arises: either through cofluctuations in the excitatory drive or normalization signal (determined by *ρ*_*N*_, *ρ*_*D*_ respectively). To employ these insights when fitting to data, we need to know how well we can infer these parameters from data. To do so, we generated synthetic neural data using realistic values for the single-neuron parameters (see subsection 2.6) and uniformly randomly sampled *ρ*_*N*_, *ρ*_*D*_ parameters in the range [-0.9,0.9]. We then assessed the quality of the maximum likelihood estimate of the parameters by calculating bootstrapped confidence intervals (with N=1000 bootstrap samples) and comparing the estimator and true values (see subsubsection 2.3.2).

First, we assessed the validity of the confidence interval by examining how well the empirical coverage probability matches the confidence level as constructed. To do so, we constructed 90% confidence intervals for the (*ρ*_*N*_, *ρ*_*D*_) parameters via bootstrap resampling. We then grouped by the ground truth *ρ* values using a sliding window and counted the proportion of cases in that bin for which the bootstrap confidence interval contains the true value. We found that for both *ρ*_*N*_ and *ρ*_*D*_ the coverage is near the nominal level: for *ρ*_*N*_ it is slightly lower (Fig. 3a left) while for *ρ*_*D*_ it is nearly equivalent (Figure 3a). This indicates that the confidence intervals constructed via the bootstrap are valid and can be used for further analysis.

Next, we directly compared the the true and inferred *ρ* values (Figure 3b). In general, the maximum likelihood estimators are largely unable to recover the true generating values (the overall Pearson correlation between the fit and true (*ρ*_*N*_, *ρ*_*D*_) = (0.48, 0.18), the mean squared error across pairs is (0.32, 0.57)), indicating that these parameters are not identifiable with the parametrization we considered (contrast tuning). Further, this analysis indicates that inference of *ρ*_*D*_ is in general more difficult than it is for *ρ*_*N*_. The lack of identifiability of these parameters is likely due to the numerous multiplicative interactions between the parameters. For instance, by looking closer at Equation 7, we can see that the contribution of the *ρ* parameters to the noise correlation is multiplied by the respective standard deviations for the numerator or denominator variables. Such interactions may make it difficult to infer the exact value of the *ρ* parameters (see subsection S7 for further discussion).

As we established the validity of the bootstrapped confidence intervals (Figure 3a), one could select pairs for which the parameter inference is accurate to the desired precision (i.e., selecting those pairs whose confidence intervals are a certain width). Additionally, one can also perform population-level analyses on the *ρ* parameters, as we demonstrate for experimental data (see subsection 3.4). Here, we introduce and validate another kind of analysis one can perform with the bootstrapped confidence intervals. Rather than the exact magnitude of the *ρ* parameters, we are often only interested in the *sign* of these parameters, as in deriving the relationship between normalization strength and noise correlations (subsection 3.1). Moreover, because all the parameters in Equation 7 are *a priori* positive besides the *ρ* parameters, we reasoned that the accuracy of sign inference might be higher. Therefore, we considered the subset of cases where the parameters are significantly different from 0, which we define as cases where the 90% confidence interval does not include 0 and the pairwise goodness of fit is greater than the independent goodness of fit (see subsubsection 2.3.1). First, for those pairs (for *ρ*_*N*_, 4598/11628 pairs were significant, for *ρ*_*D*_, 1625/11628), we see a much stronger relationship between the fit and true *ρ* values (Figure 3b, darker colors; Pearson correlation between significant fit and true (*ρ*_*N*_, *ρ*_*D*_) = (0.89, 0.57); mean squared error across pairs = (0.09,0.43)). Importantly, the plots also show that, for pairs with *ρ* parameters significantly different from 0, the sign of the inferred *ρ* parameter is very frequently equivalent to the sign of the true *ρ* parameter, in a similar proportion of cases for *ρ*_*N*_ and *ρ*_*D*_, although *ρ*_*N*_ has a much higher proportion of cases (for *ρ*_*N*_, 4509/4598 of pairs significantly different from 0 has the same sign; for *ρ*_*D*_, 1309/1625). From this, we conclude that, for these significant *ρ* parameters, the *sign* of the inferred *ρ* parameter is accurate.

In summary, our analysis indicates that it is difficult to estimate the precise value of the *ρ* parameters in general, for the stimulus parametrization considered here (i.e., the classical normalization model for contrast tuning). However, we have provided a method to calculate bootstrapped confidence intervals for the maximum likelihood estimators of the *ρ* parameters and have shown that these confidence intervals accurately represent the uncertainty around those estimates. We then demonstrated one possible use-case for these confidence intervals: for *ρ* estimates that are significantly different from 0, the *ρ* estimators are accurately able to recover the sign of the ground truth *ρ* parameters.

**Figure 3:**
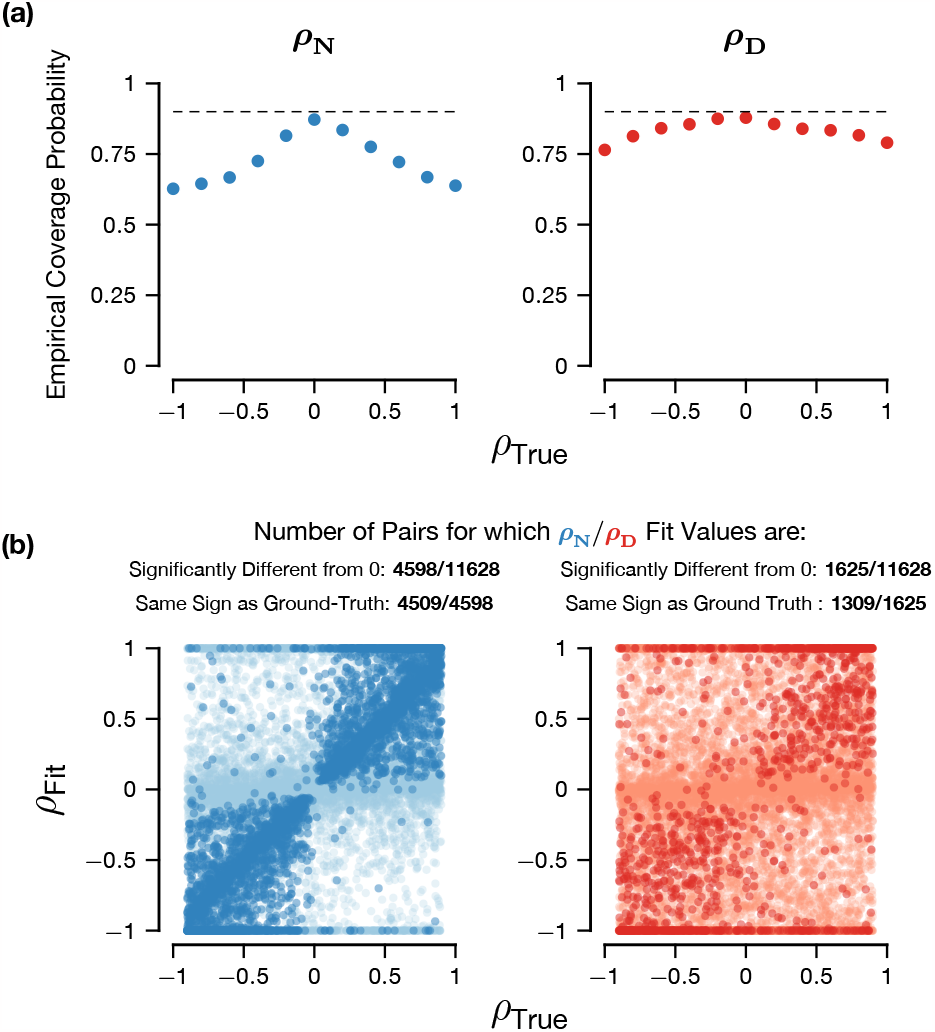
Accuracy of Inference of *ρ* Parameters. Plots were generated with 11628 synthetic parameter pairs with uniformly randomly generated *ρ*_*N*_, *ρ*_*D*_ ∈ [−0.9, 0.9], contrast levels {6.25, 12.5, 25, 50, 100}, 1000 synthetic trials and 1000 bootstrap resamples. The left column are the results of the analysis for *ρ*_*N*_, the right column for *ρ*_*D*_.(a) Empirical coverage probability for the 90% confidence intervals as a function of the ground truth *ρ* values. The dotted line indicates the nominal confidence level. Coverage probability was computed as the proportion of cases with a specified range of *ρ* values for which the 90% confidence intervals contained the true value. We used a moving window with width 0.4 and a step size of 0.2.(b) Direct comparison between the true *ρ* value and the maximum likelihood estimator for the *ρ* value. The darker colors are the pairs for which the *ρ* parameters are significantly different from 0 (i.e., the 90% confidence interval excludes 0), whereas the lighter colors are not significant (i.e., the 90% confidence interval includes 0).

### 3.3 Pairwise Model Improves Single-Trial Inference of Normalization Strength Even When Noise Correlations are Small

In past work that connected normalization to modulation of noise correlations, stimulus and experimental manipulations (e.g., contrast, attention) are used as proxies for normalization strength [50, 51, 54] because normalization strength cannot be measured directly. However, these manipulations also affect other factors that drive neural responses (as we have illustrated in subsection 3.2, Figure 3), which could confound these as measures of normalization signal. Therefore, quantitatively testing the relationship between noise correlations and normalization requires estimating the single-trial normalization strength for a pair of neurons. One of the advantages of our probabilistic, generative formulation of the pairwise RoG model (Equation 2) is that it allows us to infer the single-trial normalization strength from measured neural activity (see subsection 2.5). The independent RoG model also provides an estimate for the single-trial normalization, which is known to be a valid estimator for the ground-truth normalization strength for data generated from the independent RoG [52]. We found similar results for the pairwise RoG (not shown), so we examined how the pairwise estimates for the single-trial normalization strength compares to estimates based on the independent model.

One possibility is that the estimate derived from the pairwise model would outperform the independent model as the magnitude of noise correlations increase. However, this is not necessarily the case. Figure 4a,b demonstrates this with two example synthetic neuron pairs. Because the single-trial normalization inference depends on the single-trial neural activity, correlations between neurons will induce correlations between the inferred normalization signals even for the independent RoG (Figure 4a). However, when noise correlations are small due to cancellation between *ρ*_*N*_ and *ρ*_*D*_, the independent model will infer minimal correlation between normalization signals while the pairwise model will correctly infer that the single-trial normalization is correlated (Figure 4b).

**Figure 4:**
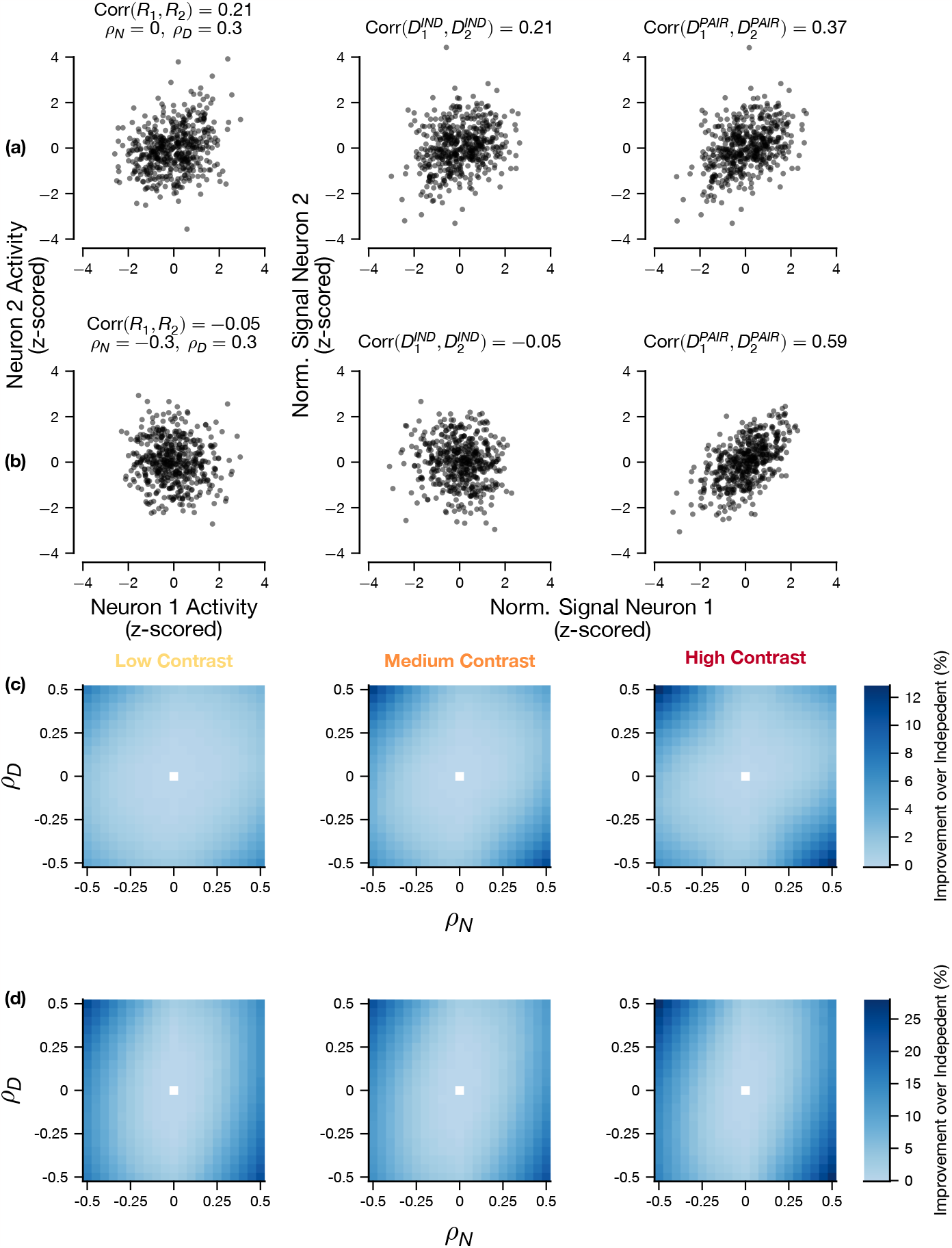
Inference of Single-Trial Normalization Depends on *ρ* Parameters and Contrast Level. (a,b) Two simulated experiments drawn from the pairwise RoG with different noise correlations arising from different underlying *ρ*_*N*_, *ρ*_*D*_ values to compare the pairwise and independent estimators of single-trial normalization with the ground truth normalization signal. (a) has overall noise correlation of 0.21 across contrasts generated with *ρ*_*N*_ = 0, *ρ*_*D*_ = 0.3, (b) has overall noise correlation of -0.05 across contrasts generated with *ρ*_*N*_ = −0.3, *ρ*_*D*_ = 0.3. Z-scoring performed across trials. Random parameters drawn from *R*^max^ ∈ [10, 100], ϵ ∈ [15, 25], α_***N***_, ***α***_***D***_ ∈ [0.1, 1], ***β***_***N***_, β_***D***_ ∈ [1, 1.5], *η* ≡ 0. Contrasts levels were {6.25, 12.5, 25, 50, 100}. (c,d) Comparison of the single-trial normalization inference in the pairwise and independent RoG models for the mean-squared error (c) and the correlation between the estimate and the true value (d), as it depends on *ρ*_*N*_, *ρ*_*D*_ and the contrast levels. Left: lowest contrast level (6.25); middle: intermediate contrast level (25); right: full contrast (100). Each bin corresponds to the median difference between the pairwise and independent models across simulated experiments. (c,d) used 11628 synthetic pairs (see subsection 2.6).

To demonstrate this principle, we computed the difference of the mean squared error (Figure 4c) and correlations (Figure 4d) between the pairwise RoG and independent RoG normalization inference (normalized with respect to the ground truth values) for many simulated pairs (11628) with systematically varying *ρ*_*N*_, *ρ*_*D*_ values. First, we found that the distinction between the two models depended on the contrast level: as contrast levels increase, the quality of the pairwise and independent estimators of normalization signal become more distinct. Second, the improvement of the pairwise RoG estimate over the independent estimate increased when the magnitude of the *ρ*_*D*_ parameter increased. Consistent with the intuition provided by Figure 4b, the largest improvement occurred when the *ρ*_*N*_ parameter had a large value that was the opposite sign of *ρ*_*D*_. The dependence on the *ρ*_*D*_ parameter reflects that the estimator for the pairwise model incorporates knowledge about correlation between the normalization signals.

In summary, this analysis shows that the pairwise model estimates of the single trial normalization can improve upon the independent model even when noise correlations are small. As the single-trial normalization estimator from the independent model was previously shown to be accurate [52], our results imply that the pairwise model estimate is also able to recover the ground-truth normalization strength. Additionally, we have outlined the conditions in which those estimates are preferable to those obtained with the independent model.

### 3.4 Pairwise Ratio of Gaussians Model Captures Correlated Variability in Mouse V1

To test how well the model captures experimental data, we applied it to calcium imaging data recorded in V1 of mice responding to sinusoidal gratings of varying contrast levels (see subsection 2.7). We analyzed neurons that were strongly driven by the visual stimuli (N=295 neurons, 5528 simultaneously recorded pairs; see subsubsection 2.7.5 for inclusion criteria). We focused on stimulus contrast tuning (Equation 8) because the formulation of the corresponding standard normalization model captures firing rate data well [35], and visual contrast affects both normalization strength and the strength of noise correlations [40].

Because the RoG framework has not been validated before on mouse V1 fluorescence data, we first applied the independent RoG and found that it provided excellent fits (average cross-validated goodness of fit 0.85, 95% c.i [0.846,0.858], 5163/5528 pairs with goodness of fit > 0.5) on par with that found in macaque V1 data recorded with electrode arrays [52], thus demonstrating that the RoG framework is flexible enough to capture datasets with different statistics. In both cases, the analysis was performed on visually responsive neurons, that therefore exhibited strong contrast tuning of the firing rate, partly explaining the high performance. However, the independent RoG could not capture correlated variability, which was prominent in the data (median noise correlation across all pairs and contrasts 0.117, c.i. [0.115, 0.120], 2781/5528 pairs had noise correlations significantly different from 0).

Therefore, we tested if the pairwise RoG could capture correlated variability in the data. Figure 5a,b demonstrates that the model can capture contrast-dependent noise correlations, both for pairs with positive (example in Figure 5a; 4327/5528 pairs) and negative median noise correlations (example in Figure 5b; 1201/5528 pairs). Importantly, even though the *ρ*_*N*_, *ρ*_*D*_ parameters were stimulus-independent, the pairwise model captured substantial changes in noise correlations with contrast for many of the pairs analyzed (2991/5528 had greater than 0.5 correlation between the observed noise correlations and model fit, across contrast levels). However, this ability to capture correlations comes at the cost of larger model complexity. To account for this, we next compared quantitatively the pairwise and independent models, using a cross-validated goodness of fit score (Equation 12). The pair-wise model slightly outperformed the independent model on average and for most pairs (median difference in goodness of fit = 0.0121, *p* < 0.001, 4123/5528 pairs with pairwise goodness of fit greater than independent), denoting that the additional free parameters are warranted. Furthermore, because the independent model is a special case of the pairwise with noise correlations fixed at zero, we found as expected that the performance difference between the two models increased for pairs of neurons with larger noise correlations (see Figure S1).

To benchmark the RoG against a widely adopted alternative model, we considered the modulated Poisson model that was previously shown to capture noise correlations in macaque V1 [66]. For application to our imaging dataset and for a fair comparison with the RoG, we used Gaussian noise instead of Poisson [73] and termed this the Modulated Gaussian (MG) model (see subsection 2.4). The example pairs demonstrate that, while in some pairs the MG can capture the modulation of noise correlations with contrast as well as the RoG (Figure 5a), it is not able to capture it in other pairs while the RoG can (Figure 5b). Across the dataset, for the majority of pairs (5238/5528), the pairwise RoG had a higher goodness of fit score (Equation 12) than the MG (Figure 5c,d, median difference between goodness of fit for RoG and MG =0.238, 95% c.i. [0.232,0.245]). These results were also largely independent of the specific preprocessing method applied to the calcium imaging data (see subsection S6 and Figure S2). Moreover, although both models capture the tuning of noise correlations with contrast level by using stimulus-independent correlation parameters, the RoG model better predicts the trend in noise correlations with contrast than the MG (median difference in correlations between model fit noise correlations and empirical noise correlations = 0.0771, 95% c.i. [0.071, 0.084]; 1450/5528 pairs had statistically significant correlation between the pairwise RoG predictions and empirical noise correlations compared to 1008/5528 for the MG). In principle the MG model’s ability to capture the modulation of noise correlations with contrast could be improved by including contrast dependence in the correlation parameters explicitly, although this would increase model complexity.

**Figure 5:**
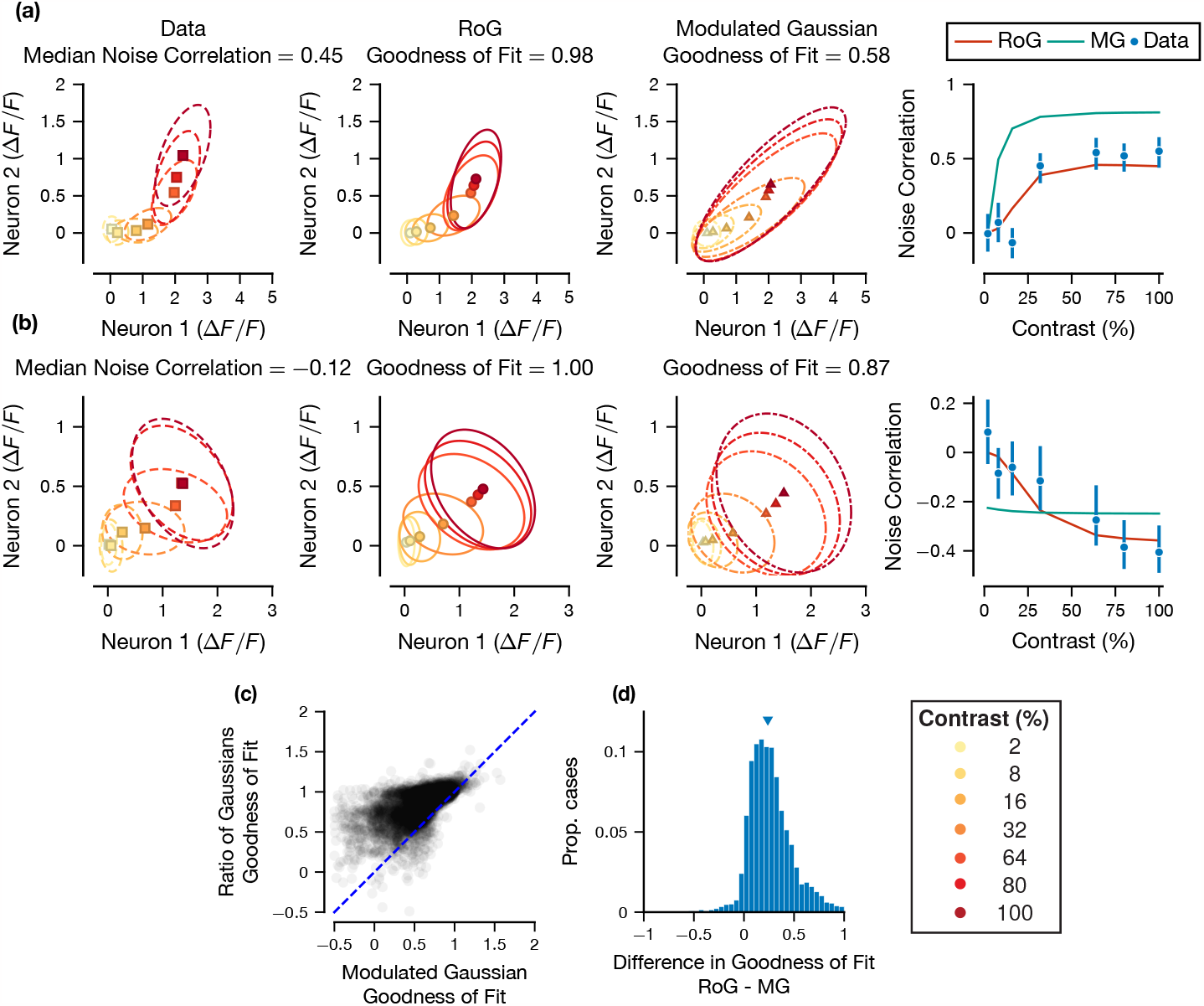
Pairwise RoG Captures Contrast-Dependent Noise Correlations in Mouse V1. (a,b) Pairwise neural responses for two example pairs of neurons in mouse V1 with positive (a) and negative (b) median noise correlations. From left to right: 1) empirical mean and covariance ellipses (∼ 1 standard deviation from the empirical mean) for pairwise responses at each contrast level; 2) the RoG model predicted means and covariance ellipses, the panel title includes the cross-validated goodness of fit score; 3) the Modulated Gaussian (MG) predicted means and covariance ellipses; 4) compares the two model fit noise correlation values (continuous lines), with the empirical values as a function of contrast (error bars are 68% bootstrapped confidence interval). Neuronal pair in (a) had 93 repeats of each stimulus contrast, pair in (b) had 68 repeats. (c) Scatter plot across all pairs of the goodness of fit score for Modulated Gaussian vs. the goodness of fit for the RoG. (d) Histogram of the difference between the scores in (c). Contrast levels are {2,8,16,32,64,80,100}.

These results demonstrate that the pairwise RoG captures a range of effects of stimulus contrast on noise correlations observed in experimental data and performs competitively against a popular alternative model that does not account for normalization explicitly.

Next, we analyzed the correlation parameters (*ρ*_*N*_, *ρ*_*D*_) in the model fit (Figure 6). We first only selected those pairs of neurons whose pairwise goodness of fit exceeded 0.5 and the independent goodness of fit measure (3920/5528 total), and we computed the bootstrapped 90% confidence interval for the (*ρ*_*N*_, *ρ*_*D*_) parameters (see subsubsection 2.3.2).

Examining the correlation parameters for all of the pairs meeting the goodness of fit criteria (Figure 6a,b outlined), we see a significant bias towards positive values (median *ρ*_*N*_, *ρ*_*D*_ parameter values = 0.84, 1). This is partially due to the large number of cases in which the fit parameters were exactly equal to ±1 (for *ρ*_*N*_, *ρ*_*D*_, 1442 and 2700 fit values were ±1). However, even when excluding these pairs, the trend within the population is still towards positive fit *ρ* values (median fit *ρ*_*N*_, *ρ*_*D*_ values excluding extreme pairs is 0.29, 0.22. This analysis suggests that these signals are, on aggregate, shared among the population recorded; in particular, this suggests that normalization is typically shared between the pairs recorded.

As a complementary analysis, we then focused on the cases where the parameters were assessed to be significantly different from zero (1270/3920 for *ρ*_*N*_, 192/3920 for *ρ*_*D*_) (Figure 6a,b filled). The proportion of pairs for which the estimated *ρ* parameters are significantly different from 0 is similar to the synthetic data (see subsection 3.2). For these pairs that were significant, we found that nearly all inferred *ρ*_*N*_ (1239) and *ρ*_*D*_ (191) parameters were positive (see Figure 6), suggesting that normalization signals are generally shared for these pairs of neurons.

In summary, our results demonstrate a new approach to quantify how strongly normalization signals are shared between neurons, and to explain the diverse effects of normalization on noise correlations.

**Figure 6:**
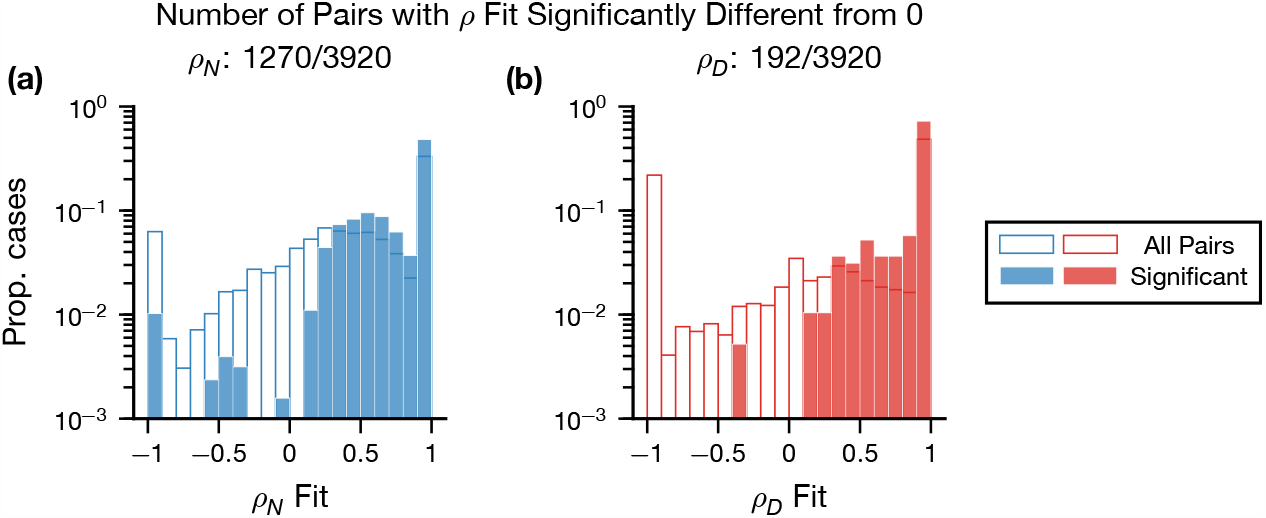
Inferred *ρ*_*N*_, *ρ*_*D*_ are Positive in Mouse V1. Histograms comparing the inferred *ρ*_*N*_ (a) and *ρ*_*D*_ (b) values for all neuronal pairs meeting our goodness of fit criteria (outlined) and the subset of those pairs significantly different from zero with 90% confidence (filled). The histograms for all pairs and pairs significantly different from 0 are normalized separately.

## 4 Discussion

We introduced a stochastic model of divisive normalization, the pairwise RoG, to characterize the trial-to-trial covariability between cortical neurons (i.e., noise correlations). The model provides excellent fits to calcium imaging recordings from mouse V1, capturing diverse effects of stimulus contrast and normalization strength on noise correlations (Figs. 5, 6). We demonstrated that the effect of normalization on noise correlations differs depending on the sources of the variability, and that the model can accommodate both increases and decreases in noise correlations with normalization (Figure 2) as past experiments had suggested. We then investigated the accuracy of inference of a key model parameter, which determines whether normalization is shared between neurons, and we provided a procedure for quantifying the uncertainty of this inference using bootstrapping (Figure 3). Lastly, we derived a Bayesian estimator for the single-trial normalization signals of simultaneously recorded pairs. Surprisingly, this estimator can be more accurate than the estimator based on the model that ignores noise correlations (the independent RoG) even when noise correlations are negligible (Figure 4).

As a descriptive, data analytic tool, our modeling framework complements normative and mechanistic theories of neural population variability. For instance, normative probabilistic accounts of sensory processing have suggested that divisive normalization may play a role in the inference of perceptual variables by modulating neural representations of uncertainty [15, 72, 74, 83–85, 132]. Similarly, normalization could play a key role in multisensory cue combination [38, 86, 87]. However, the posited effect of normalization on covariability has not been tested quantitatively, as normalization signals are often not measurable. The pairwise RoG will allow researchers to test these hypotheses by providing a means with which to estimate normalization signals from neural data and relate these to measures of neural covariability. In circuit-based models of neural dynamics such as the stabilized supralinear network [24] and the ORGaNICs architecture [88], the normalization computation emerges naturally from the network dynamics [25] and shapes the structure of stimulus-dependent noise correlations [89]. By quantifying the parametric relation between normalization and covariability, our descriptive tool will enable mapping those parameters onto the different circuit motifs and cell types posited by these network models.

When comparing the RoG to the Modulated Gaussian model (see subsection 2.4), we found that the RoG had better performance for the majority of pairs (Figure 5c). We chose to adapt the modulated Poisson model [66] as a comparison to the RoG because it was shown to successfully capture noise correlations in recordings from macaque V1. Moreover, this model belongs to the class of Generalized Linear Models, which are among the most widely used encoding models for neural activity [26, 90]. There are numerous alternative descriptive models of correlated neural population activity, among the most popular of these being latent variable models (LVMs), in which population-wide activity arises from interactions between a small set of unobserved variables [27, 32, 91–94]. This effectively partitions the population noise covariance into underlying causes (i.e., latents) that are responsible for coordinating neural responses, which resembles our attribution of noise correlations to either shared input drive or shared normalization pools (Figure 2). The RoG, on the other hand, is a pairwise model that seeks to explicitly characterize neural interactions through divisive normalization, which cannot be done with any existing LVMs; integrating normalization into the LVM framework is an important future extension of our model. One benefit of our current approach is that the RoG can be applied to any scenario in which two or more neurons are simultaneously recorded. LVMs can only be applied to relatively large populations of simultaneously recorded neurons to estimate the globally shared latent factors. This is not always feasible for regions of the brain that are difficult to record from or using techniques such as intracellular voltage recordings [95, 131]. The downside of a method such as the RoG is scalability to large populations, as the model parameters must be optimized for each recorded pair, which can be computationally expensive for modern datasets with thousands of neurons [96]. Nonetheless, we were able to fit the RoG to data across multiple different preprocessing methods (∼ 90000 pairs total) in a reasonable time (∼ 27 hours running in parallel on a 28-core server without GPU acceleration), suggesting that it is not entirely impractical to use the pairwise RoG on a large dataset.

Three models have directly studied the relationship between normalization and across trial covariability [50,51,97]. Tripp’s [50] simulation work on velocity tuning in the medial temporal cortex (MT) consistently predicted that normalization would decorrelate neural responses. However, we found that noise correlations could also increase with normalization. This is because Tripp modeled correlations to solely arise from tuning similarity between neurons. Conversely, in the RoG framework, noise correlations originate from input correlations *ρ*_*N*_ and correlations between normalization signals *ρ*_*D*_. Our model then offers more flexibility than Tripp’s by allowing relationships between normalization and correlation to depend on the sources of correlations. Verhoef and Maunsell [51] investigated the effect of attention on noise correlations by using a recurrent network implementation of the normalization model of attention [48]. They describe multiple different patterns of the effect of normalization on noise correlations depending on tuning similarity between a pair of neurons and where attention is directed. Our model does not currently account for the effect of attention, but this would be possible by adapting the standard normalization model of attention which would require an additional parameter for the attentional gain. These prior two models are also primarily simulation based, while our model is meant to be data analytic. Lastly, Ruff and Cohen [97] proposed a normalization model to explain how attention increases correlations between V1 and MT neurons [54]. They modeled the trial-averaged MT neural responses as a function of trial-averaged responses of pools of V1 neurons. After fitting the parameters, single-trial MT responses were predicted by feeding the pooled single-trial V1 responses into the equation. By construction, variability in predicted MT neural responses only arises from variability in the V1 neural responses, which only occur in the numerator of their normalization model. Our model also allows for variability in the denominator of the normalization equation and therefore their model can be seen as a special case of the pairwise RoG.

An important limitation of our model is that the correlation parameters (*ρ*_*N*_, *ρ*_*D*_) are not identifiable (Figure 3), meaning the model parametrization is such that multiple different parameter sets result in equivalent models (e.g., equivalent likelihoods and moments). This is a common issue when using complex nonlinear models as proposed here [98]: in our model, this is due to multiplicative interactions between model parameters. Improving parameter estimation will require better constraints on model parameters, alternative optimization algorithms, or different objective functions (see subsection S7 for further discussion). Future extensions of the model to population level interactions through latent variable models offer another avenue to improve parameter estimation: the variability of population activity is often low-dimensional, which could naturally impose parameter constraints. Nonetheless, we developed a method to calculate confidence intervals for the estimates of the *ρ* parameters, which can be used to select pairs for which the estimates’ uncertainty is less than a desired level. As an example application of this approach, we have demonstrated that the confidence intervals can be used to determine when the sign of those parameters, which is an important factor in controlling noise correlations (Figure 2), can be recovered accurately. We showed that we were accurately able to recover the sign of the correlation in synthetic datasets when the bootstrap confidence interval for the parameter of interest excluded 0. In the V1 dataset analyzed, we found that (1239,192)/3920 pairs meet this criterion for *ρ*_*N*_, *ρ*_*D*_ respectively, and that the vast majority of those pairs had positive *ρ*_*N*_, *ρ*_*D*_. Although this is a minority of cases, it demonstrates that typical datasets with existing recording technologies could nonetheless provide sufficient power for studies that focus on the *ρ* parameters values. It will be important in future work to understand which experimental conditions would maximize the yield of pairs with accurate estimates of the *ρ* parameters.

We chose in this study to primarily analyze the normalized fluorescence traces (Δ*F*/*F*) rather than using deconvolution or spike inference methods (see [99–101] for a review). Deconvolution methods were developed in part due to the slow temporal dynamics of the calcium indicators relative to membrane potentials generating spiking activity [102, 103]. Deconvolution and other spike inference techniques attempt to mitigate this limitation for analyses that depend on more exact measures of spike timing, and developers note these methods should be avoided when temporal information is not relevant and the raw calcium traces provide “sufficient information” [100]. Because of the construction of the contrast detection task (see subsection 2.7) and the temporally invariant nature of contrast responses in V1 [104], the analysis of the dataset presented here does not require precise temporal information, so the use of normalized fluorescence traces was sufficient. Additionally, deconvolution changes the statistics of the data greatly, such as altering the distribution of noise correlations and increasing the sparsity of the fluorescence signal [105]. One recent work attempted to account for these differences by using more appropriate probabilistic models [106] but does not currently model noise correlations. On the other hand, calcium fluorescence is an indirect measure of neuronal communication and coding, being related to the underlying action potentials through a complex generative model [107,108]. As such, it might be inappropriate or insufficient to apply an encoding model directly to the Δ*F*/*F* traces, as we have done here. To address this concern, we additionally analyzed deconvolved traces using two variants of the OASIS method [109]: unconstrained OASIS as found in suite2p [80], or OASIS with an ℓ_1_ sparsity constraint as in [110]. As expected, the deconvolution techniques significantly altered the distribution of noise correlations, but the results of our analysis of these deconvolved data was qualitatively in-line with the results obtained on the raw calcium traces (see subsection S6).

The generality of the modeling framework presented here leaves room for future expansion. One such direction would be to increase the dimensionality to model correlations among a neural population. This would require more correlation parameters, which could make the model more difficult to fit to data. However, reasoning that population variability is low-dimensional [81, 111–115], it is likely this issue could be circumvented by applying dimensionality reduction techniques within the model or by allowing the sharing of correlation parameters across a neural population. Another interesting application of this model would look directly at the effects of normalization on information transmission and representation. The relationship between noise correlations and the amount of information that can be represented by a neural population has been widely discussed [7–9, 11, 12, 116]. Moreover, some experimental and theoretical work has connected modulations of information in neural populations with computations that have been modeled with normalization models, such as surround suppression and attention [43, 50, 117, 118]. Our model could be modified to investigate this connection and further illuminate the effects of normalization on information transmission.

## Acknowledgement

We thank members of the Kohn and Coen-Cagli laboratories for feedback on the manuscript. We also thank Daniel Quintana for help with animal training and Kenny Ye for advice on statistical analyses.

This work was supported by the National Institutes of Health (NIH) Grant R01-EY030578 (R.C.C.), NIH Grant RF1-DA056400 (R.C.C.), NIH Grant UF1-NS107574 (H.A.), NIH Grant U19-NS107613 (H.A.), NIH Grant RF1-RF1-MH120680 (H.A.), NIH Grant RO1-EY023756 (H.A.), New York Stem Cell Foundation (H.A.), National Science Foundation Graduate Research Fellowship (DGE 1752814 (H.A.B.)).

---

**S1 Text Derivation of Relationship Between Mean Normalization Strength and Noise Correlations**. Mathematical derivations relating noise correlations and normalization in the model.

**S2 Text Derivation of Negative Log-Likelihood**. Details of the calculation of the negative-log likelihood for the Ratio of Gaussians distribution.

**S3 Text Derivation of Moments for the Generalized Model**. Moments of the Ratio of Gaussians distribution for the general case of cross-correlations between numerator and denominator.

**S4 Text Negative Log-Posterior for Inference of Single-Trial Normalization Strength**. Expands on subsection 2.5, Eq. 14, showing the coefficient expressions.

**S5 Text Pairwise Model Outperforms the Independent Model in Simulations and V1 data When Noise Correlations are Large**. Comparison of pairwise and independent Ratio of Gaussians model goodness of fits.

**S1 Figure Improvement in Fit of the Pairwise RoG over the Independent RoG Depends on the Magnitude of the Noise Correlations**. (a) Scatter plot demonstrating the dependence of the difference in fit quality on the median noise correlation (across contrasts). (b) Mean percent difference when binning by the magnitude of the median noise correlations. Data were generated as described in Fig. 3. (c) Scatter plot of the difference in goodness of fit score of the Pairwise and Independent RoG when applied to calcium imaging data (see subsection 3.4). (d) Mean difference in goodness of fit between Pairwise and Independent RoG binning by the magnitude of the median noise correlations in calcium imaging data. Error bars are bootstrapped 95% confidence intervals.

**S6 Text Analysis of Deconvolved Imaging Data**. Examining model performance on deconvolved fluorescence traces.

**S2 Figure Comparison of Raw and Deconvolved Data**. (a) Tuning of noise correlations with contrasts across all recorded pairs. The error bars are the standard error of the mean. (b) Histogram of noise correlations across all contrast levels. (c-f) Goodness of fit comparison for the RoG and the MG models, similar to Fig. 5c,d, for suite2p (c,d) and ℓ_1_ OASIS (e,f)

**S7 Text Further Disccusion of Parameter Identifiability**. Additional considerations for *ρ* parameter estimation from data.

**S3 Figure Relationship Between Noise Correlations and Denominator Strength**. Expands upon Fig. 2 to include cases where (*ρ*_*N*_, *ρ*_*D*_) can be negative or have opposite signs. Figure was created using the exact same method and synthetic dataset as Fig. 2: see the caption in the main text for details.

**S4 Figure Relationship Between Noise Correlations and Ratio of Gaussians Parameters**. Noise correlations in the model (Eq. 7) can be modulated by stimulus strength (i.e. contrast), the correlation parameters of the model (*ρ*_*N*_, *ρ*_*D*_) and the parameters of the normalization model, in this case 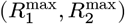and (ϵ_1_, ϵ_2_) (see 2). To understand these effects in isolation, we looked at how Eq. 7 changed with respect to each parameter, while keeping the other parameters constant. We illustrate with noise correlations that increase with contrast (a1-e1), and correlations that decrease with contrast (a2-e2). (a) Dependence of noise correlations on contrast. Three contrast levels that are fixed in the other panels are shown. (b) Dependence of noise correlations on *ρ*_*N*_. (c) Dependence of noise correlations on *ρ*_*D*_. (d) Dependence of noise correlations on 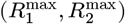, shown as a contour plot with the shade of color indicating noise correlation level. Different colors indicate different contrast levels as shown in the legend. (e) Dependence of noise correlations on (ϵ_1_, ϵ_2_). (a1-e1) uses the following parameters (when not fixed): 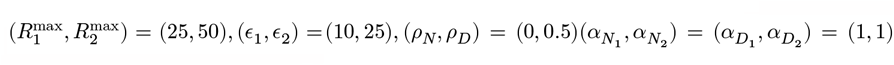, 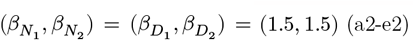 uses the same parameters except with (*ρ*_*N*_, *ρ*_*D*_) = (0.5, 0). Contrast levels were {1,…,100}.

## Supporting Information

### S1 Derivation of Relationship Between Mean Normalization Strength and Noise Correlations

We use the same notation as in subsection 2.1 (see Equation 7). The true characterization of how changes in the mean normalization strengths 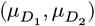 cause changes in *ρ*_*NC*_ ≔ Corr(*R*_1_, *R*_2_) (Equation 7) would be to consider the directional derivative ∇ρ_*NC*_ ·(1, 1). However, it will suffice to consider the partial derivative with respect to either 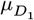 or 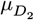. We note a few identities which are easy to verify:

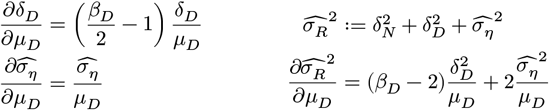

Then considering 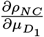, we compute using the chain rule:

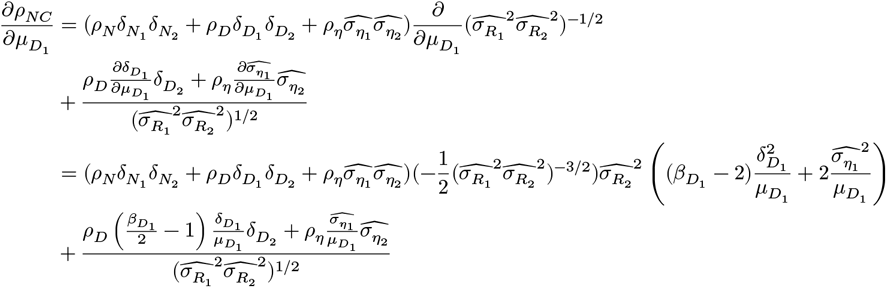

Simplifying, we get the equation:

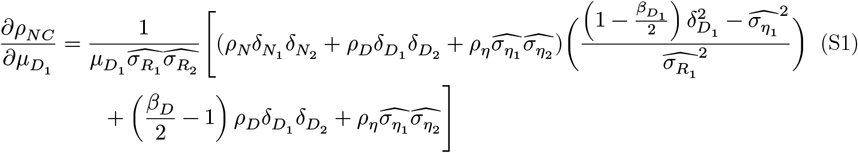

To reproduce the patterns noted in Figure 2a, we first set *η* ≡ 0 and *ρ*_*N*_ = 0, *ρ*_*D*_ > 0.

Then

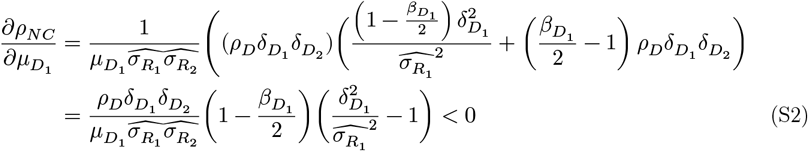

Because 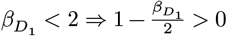. Moreover, 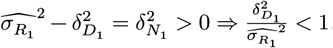. Thus, *ρ*_*NC*_ decreases as a function of normalization strength.

Similarly, when *η* ≡ 0, *ρ*_*D*_ = 0, *ρ*_*N*_ > 0 (Figure 2b), then

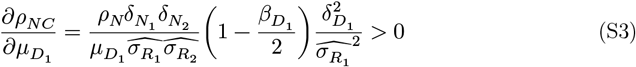

Demonstrating that *ρ*_*NC*_ increases with normalization in this case.

When *ρ*_*N*_, *ρ*_*D*_ are the opposite signs, the inequalities in (S2) and (S3) reverse and the relationship between *ρ*_*NC*_ and normalization switch to be increasing and decreasing, respectively.

Next, when *η* ≡ 0 and *ρ*_*N*_ = − *ρ*_*D*_ = *ρ*, (S1) reduces to:

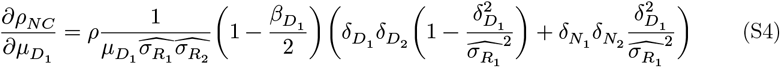

Thus, when 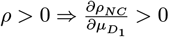. In fact, we can use (Equation S4) along with (Equation S2, S3) to generalize to all cases where *ρ*_*N*_, *ρ*_*D*_ have opposite signs. As an example, suppose *ρ*_*D*_ < 0 and *ρ*_*N*_ = |ρ_*D*_| + *C* > 0, *C* > 0. Then we can decompose 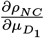 as a combination of (S4, with *ρ* = |ρ_*D*_|) and (S3, with *ρ*_*N*_ = *C* > 0). Since both these terms are positive, we conclude that *ρ*_*NC*_ increases with normalization when *ρ*_*N*_ > 0, *ρ*_*D*_ < 0, *ρ*_*N*_ > |*ρ*_*D*_|.

Lastly, we consider when *η* ≢ 0. The nature of (S1) prevents us from making any conclusions in general, as the sign of 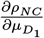 will depend on the magnitudes of the various terms, similar to when *ρ*_*N*_, *ρ*_*D*_ > 0. To get some understanding of the behavior of *ρNC*, let us assume that *ρ*_*η*_ = 0, *ρ*_*D*_ = 0, a generalization of (S3). For these assumptions, we have

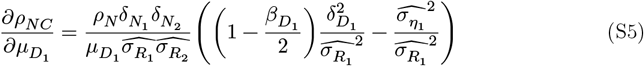

Here, if 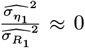 (i.e.,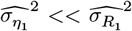), then (S5) should be approximately equal to (S3). This gives the heuristic that if the residual variance is much less than the variance of the neural response, then the presence of this noise should have little effect on these derived relationships.

### S2 Derivation of Negative Log-Likelihood for the Model

The model assumptions that the data is distributed according to a bivariate Gaussian distribution with mean and covariance given by the RoG approximation (4,5). From this, we calculate the negative log-likelihood:

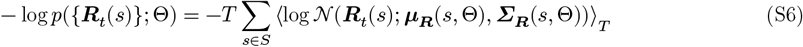

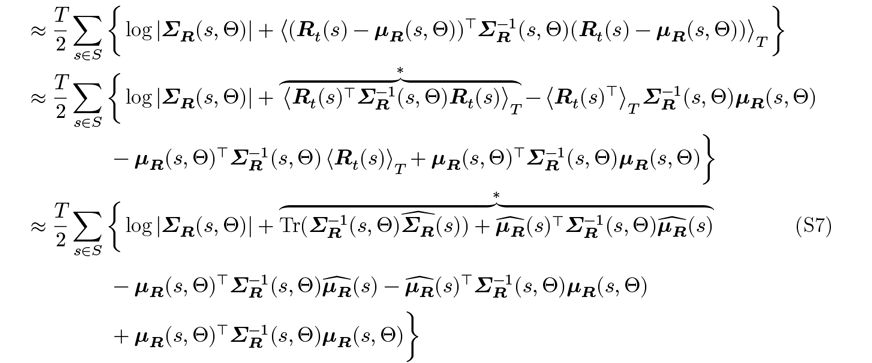

The terms marked by * are equal according to:

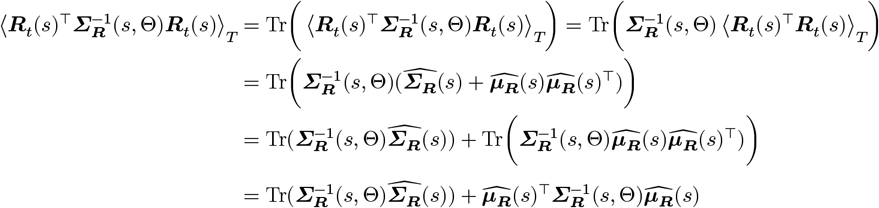

Using the fact that Tr(*AB*) = Tr(*BA*) and *c* = Tr(*c*) for *A, B* matrices and *c* a scalar. Simplifying (S7) produces (11).

### S3 Derivation of Moments for the Generalized Model

In subsection 2.1 and throughout this paper, we assumed that the numerator and denominator variables are uncorrelated, allowing us to consider two separate variables ***N***_*t*_, ***D***_*t*_ to be bivariate Gaussians. In the generalized form of the pairwise RoG generative model, we consider (***N***_*t*_, ***D***_*t*_) to be distributed according to a four-dimensional Gaussian (using the same notation as in subsection 2.1):

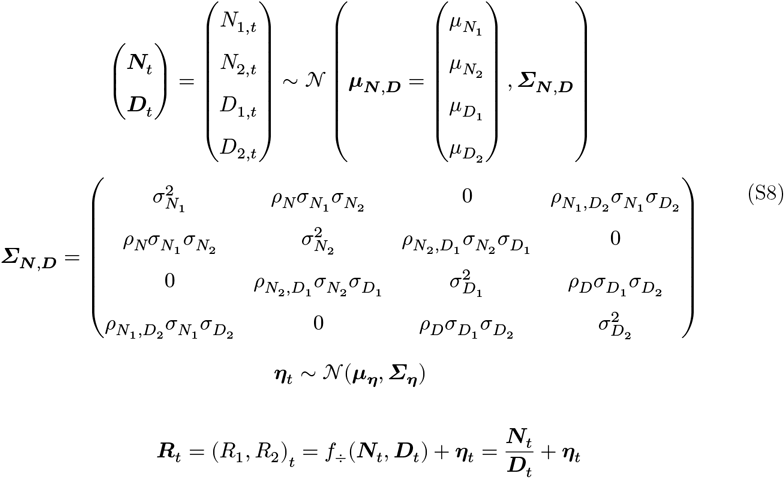

We have assumed that correlations within the equivalent independent RoG model (i.e., the correlations between *N*_1_, *D*_1_ and between *N*_2_, *D*_2_), based on our prior work that showed overfitting when including these correlations [52].

Then, we consider the Taylor expansion around μ_***N***,***D***_, as in the main text and discarding higher order terms. The mean of the ratio distribution is the same as previously derived (Equation 4). However, the derivation for the covariance is slightly different:

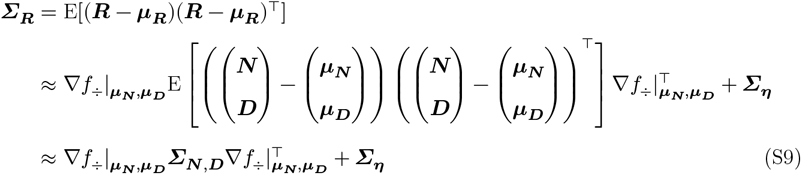

Note that, instead of the block diagonal matrix used in Equation 5, we use the full covariance matrix defined in Equation S8. The formula for the variance is the same as previously derived. For the covariance and correlation, we have:

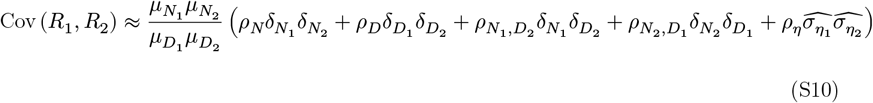

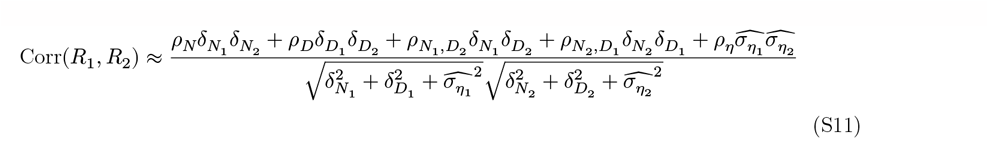

This adds two additional parameters 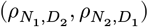 that need to be optimized using the negative log-likelihood (Equation 11). We note that the accompanying code toolbox includes these parameters.

### S4 Negative Log-Posterior for Inference of Single-Trial Normalization Strength

Using Bayes’ theorem and the change of variables formula, we compute the negative log of the probability distribution of the single-trial normalization signals (***D***_*t*_) given the observed neural activity during that trial (***R***_*t*_) (14) as:

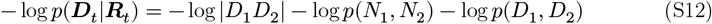

We use the generative model (2) and simplify to obtain the following expression:

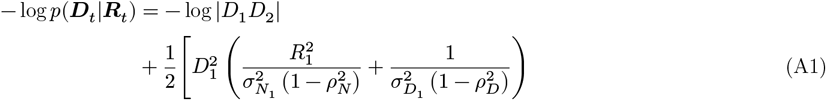

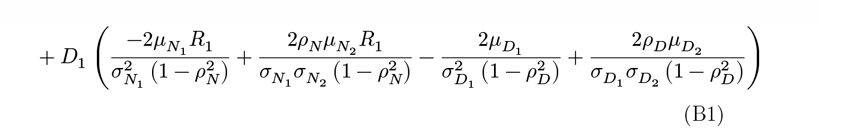

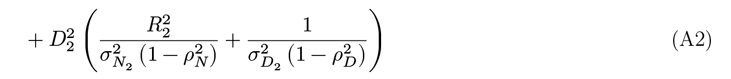

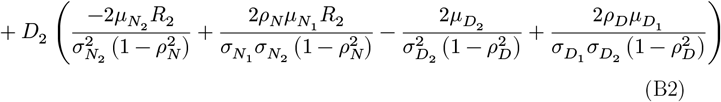

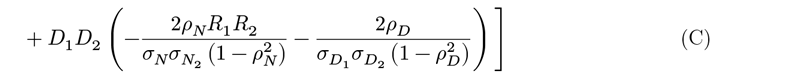

Substituting the coefficients on the right, we can write this equation as:

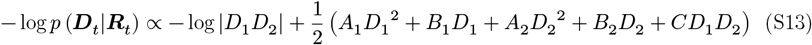

With the assumption that *D*_1_, *D*_2_ > 0 (which is expected since these are supposed to the sums of neural activity), we take the partial derivatives of (S13) to obtain the polynomials (15).

### S5 Pairwise Model Outperforms the Independent Model in Simulations and V1 data When Noise Correlations are Large

Our previous work [52] demonstrated that the independent RoG model provided an excellent fit to single-neuron data. Moreover, by construction, the independent and pairwise model best fit parameters are identical except for the correlation parameters (the *ρ* parameters, see subsection 2.3). Thus, we expected any differences in the fit quality (as measured by the negative log-likelihood using the best fit parameters) between the independent and pairwise RoG models to be minimal for pairs of neurons with noise correlations close to zero, and that the pairwise RoG would outperform the independent RoG for pairs of neurons with prominent correlated variability.

To test these predictions, we simulated pairs of neurons with parameters selected as described in Methods (see subsection 2.6) and compared the negative log-likelihood values for the pairwise and independent model (a lower negative log-likelihood denotes better goodness of fit). For these simulations, we used 6 stimulus contrasts ([6.25, 12.5, 25, 50, 100]) and 1000 trials. We used cross-validation to account for the extra free parameters of the pairwise model (details in subsection 2.3). We found that 5443/11628 of the pairs were better captured by the pairwise model, and on average the pairwise model slightly outperformed the independent model (mean percent difference = 0.964, p<0.01). We studied how the difference of likelihoods depends on the noise correlation of each pair (median noise correlation across contrasts; Figure S1a). We found that, for most pairs of neurons with magnitude of noise correlations greater than 0.1, the pairwise model outperformed the independent model (3411/4040 pairs). To quantify the dependence of this model improvement on noise correlation strength, we binned simulated pairs of neurons by the magnitude of noise correlations and computed the average difference of negative log-likelihoods for the pairwise and independent models (Figure S1b). This demonstrates that, as the magnitude of the noise correlations increases, the pairwise model provides an increasingly better fit to the data over the independent model.

We further validated this synthetic result using the calcium fluorescence data and obtained similar results (Figure S1c,d; see subsection 2.7).

### S6 Analysis of Deconvolved Imaging Data

In the main paper, we chose to analyze the raw normalized fluorescence traces (see subsection 2.7 for details of data analysis and section 4 for our rationale). However, as deconvolution of calcium traces are widely used in the literature, we also performed our analysis on the data processed with OASIS deconvolution [109]. We analyzed the data using two different constraints during the spike inference: 1) only constraining the resulting deconvolved traces to be positive (*suite2p* [80], and 2) constraining the deconvolved traces to be sparse (ℓ_1_ *OASIS*) [110]. Using deconvolution also increased the number of neurons that were assessed to be visually responsive: from the raw fluorescence traces, we assessed that 295 neurons were visually responsive (see subsection 2.7), while for suite2p there were 688 neurons and for ℓ_1_ OASIS there were 856 neurons. For comparison of the deconvolutional methods to the normalized fluorescence traces, we only analyze neurons and neuronal pairs that were considered in the main text.

First, we examined how deconvolution would change the statistics of the neuronal data, in particular how deconvolution would alter the distribution of noise correlations. From Figure S2a, we see that, on average, the changes of noise correlations with contrast are similar across each of the three preprocessing methods. The main difference among the three methods is the magnitude of the noise correlations. On average, noise correlations appear to increase with contrast for all three preprocessing methods except for a decrease in correlation strength for 80% contrast level. One possible reason for this decrease is that, for two of the recording sessions animals were not presented with a grating with this contrast level, leading to less data relative to the other contrast levels. The distributions of noise correlations are not identical, however, as seen in Figure S2b. While the normalized traces and suite2p deconvolution appear very similar, the ℓ_1_ OASIS deconvolved traces have more peaked noise correlations near 0 and skewed towards negative noise correlations. This difference in distribution may be due to the sparsity constraint inherent to this method, which would decrease the overall neuronal activity relative to the normalized traces and suite2p.

Next, we looked at how well the RoG and the Modulated Gaussian (MG) model fit to the deconvolved data (Figure S2c; see subsection 2.4 for description of the model comparison and 3.4 for the corresponding analysis using the normalized calcium traces). We see that, for suite2p, there is only a marginal improvement in goodness of fit for the RoG relative to the MG, while for ℓ_1_ OASIS the RoG outperformed the MG, as was found in the normalized calcium traces (Figure 5c,d).

### S7 Further Discussion of Parameter Identifiability

In sections 3.2 and 4 we investigated and discussed the accuracy of the estimation of model parameters, namely the *ρ* parameters. We hypothesize that the inability for the optimization to capture the magnitude of these parameters is due to the parametrization of the variance of the numerators and denominators and how they interact with the *ρ* parameters in the equation for the noise correlations (Equation 7). In the ensuing discussion, we focus on one possible multiplicative interaction between model parameters that may impede exact inference of the *ρ* parameters to give intuition about the problem. In full generality, there are many multiplicative interactions involving the *ρ* parameters (Equation 7), including the mean of the numerator and/or denominator variables raised to a power (depending on the parameter β) and the variances of the neural responses, which are functions of the means and variances for the numerators and denominators (Equation 6). Additionally, as the means of the the numerators and denominators involve further parameters describing the base normalization model (e.g., the contrast response function, Equation 8), it would be difficult to fully describe all the possible multiplicative interactions involving the *ρ* parameters and other model parameters that are fit to data.

As a simplified example of this phenomenon, consider Equation 7 with *ρ*_*D*_ = 0, *ρ*_*η*_ = 0, all β = 2,, and 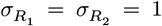(this can be achieved manually tuning 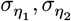), then we have that 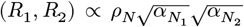. This effectively partitions a single number (the noise correlation) into three different contributions, which makes it difficult to correctly infer the magnitude of each parameter. In the two-step optimization procedure we use in this paper (see subsection 2.3), these parameters are estimated during the first phase in which correlations are ignored. If these parameters are incorrectly estimated to be at the extremes of the broad parameter bounds, when the *ρ* parameters are estimated in the second phase of the optimization, they will be forced to the extremes of their ranges to compensate. Using our previous example, suppose that α are all (incorrectly) estimated to be 0.1. If the noise correlation is measured to be 0.1, this will force *ρ*_*N*_ = 1. This is an extreme example, but is illustrative of the issue with estimating these parameters. This problem becomes compounded by the presence of the other *ρ* parameters needing to be optimized.

One solution to this problem would be to put tighter constraints on the model parameters, primarily the α, β parameters as the parameters of the base normalization model (e.g., the contrast response function, Equation 8) are more explicitly constrained by the data. The bound constraints for α, β are hyperparameters that may need to be tuned explicitly. It is *a priori* difficult to constrain the α, β parameters as they represent the mean-variance dependence for internal variables of the model (numerator and denominator) which are not measurable. We currently have a very broad range of α ∈ [0.1, 20]; the range of β ∈ [1, 2] is more restricted under the assumption that the numerator and denominator variables should overdispersed, as is often found in neuronal data ([66] but see [?,?]). One possible additional constraint would be to replace the eight α, β parameters with two global parameters, α = *a*, β = *b* for some *a, b*, or have separate parameters for the numerators and denominators (α_***N***_ = *a*_*N*_, β_***N***_ = *b*_*N*_ and similarly for the denominator). These parameters could also be optimized, or they could be fixed to a specific value; for instance, by setting α = 1, which would match the mean-variance relationship found in some neural recordings. A related approach would be to constrain *ρ*_*N*_ using measurements of mean firing rate tuning similarity (see end of subsection 2.1). Complementary to further constraining the model by directly changing the parameter bounds or dependence among variables discussed above, we can explore global optimization algorithms to improve parameter estimates or modified objective functions, namely by imposing some sort of regularization on the parameters.

**Figure S1:**
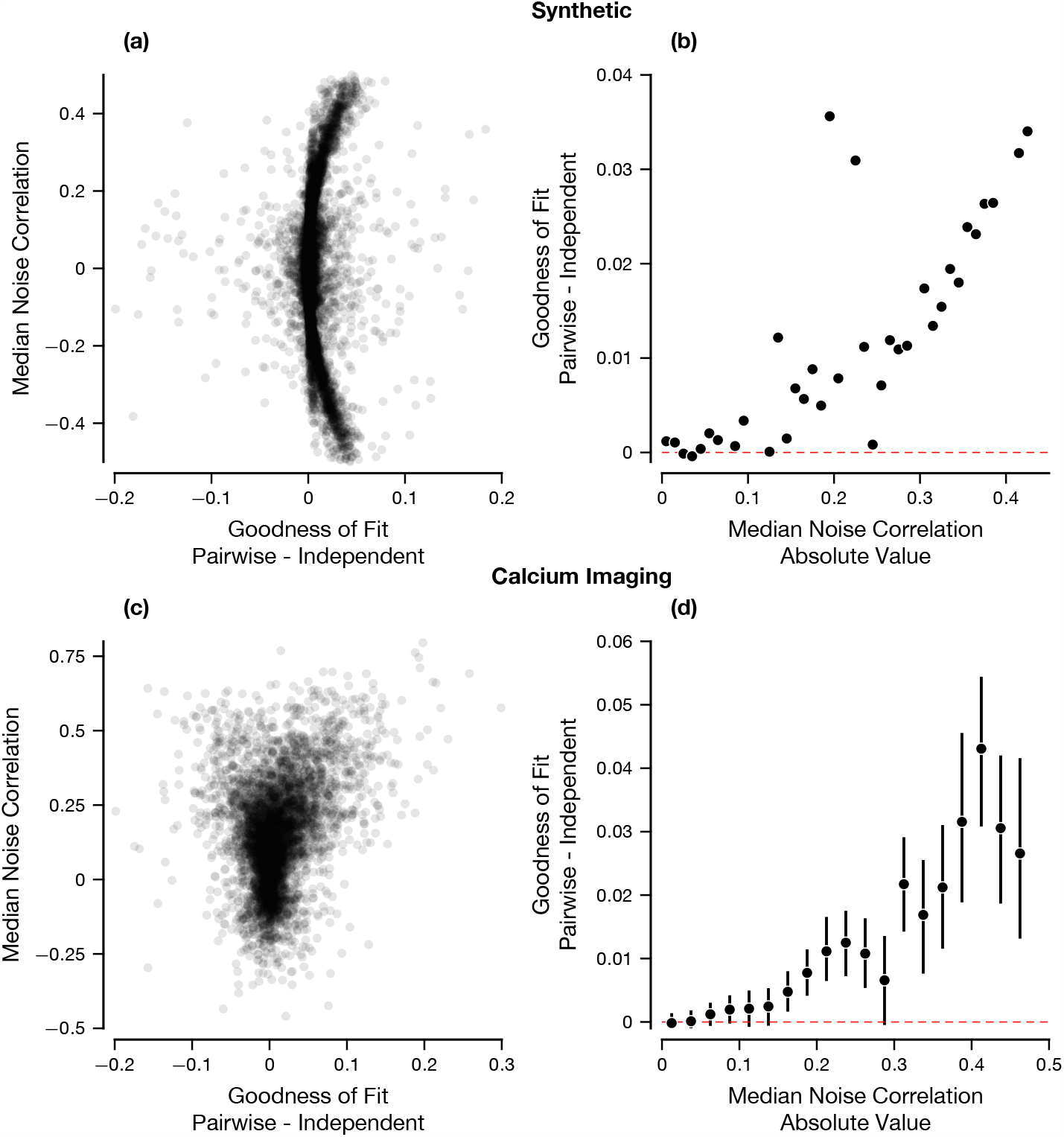
Improvement in Fit of the Pairwise RoG over the Independent RoG Depends on the Magnitude of the Noise Correlations. (a) Scatter plot demonstrating the dependence of the difference in fit quality on the median noise correlation (across contrasts). (b) Mean percent difference when binning by the magnitude of the median noise correlations. Data were generated as described in Figure 3. (c) Scatter plot of the difference in goodness of fit score of the Pairwise and Independent RoG when applied to calcium imaging data (see subsection 3.4). (d) Mean difference in goodness of fit between Pairwise and Independent RoG binning by the magnitude of the median noise correlations in calcium imaging data. Error bars are bootstrapped 95% confidence intervals.

**Figure S2:**
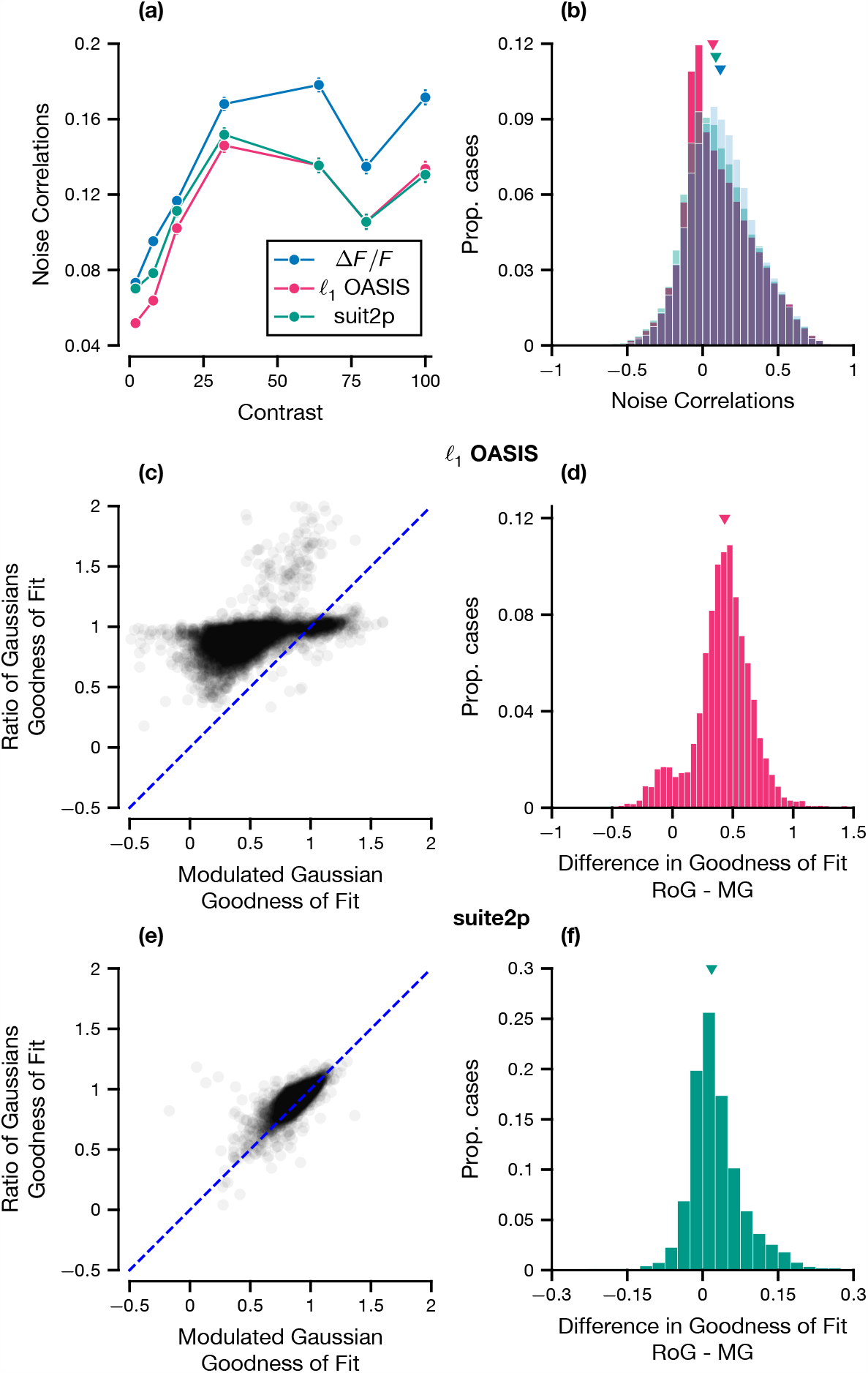
Comparison of Raw and Deconvolved Data. (a) Tuning of noise correlations with contrasts across all recorded pairs. The error bars are the standard error of the mean. (b) Histogram of noise correlations across all contrast levels. (c-f) Goodness of fit comparison for the RoG and the MG models, like Figure 5c,d, for suite2p (c,d) and ℓ_1_ OASIS (e,f)

**Figure S3:**
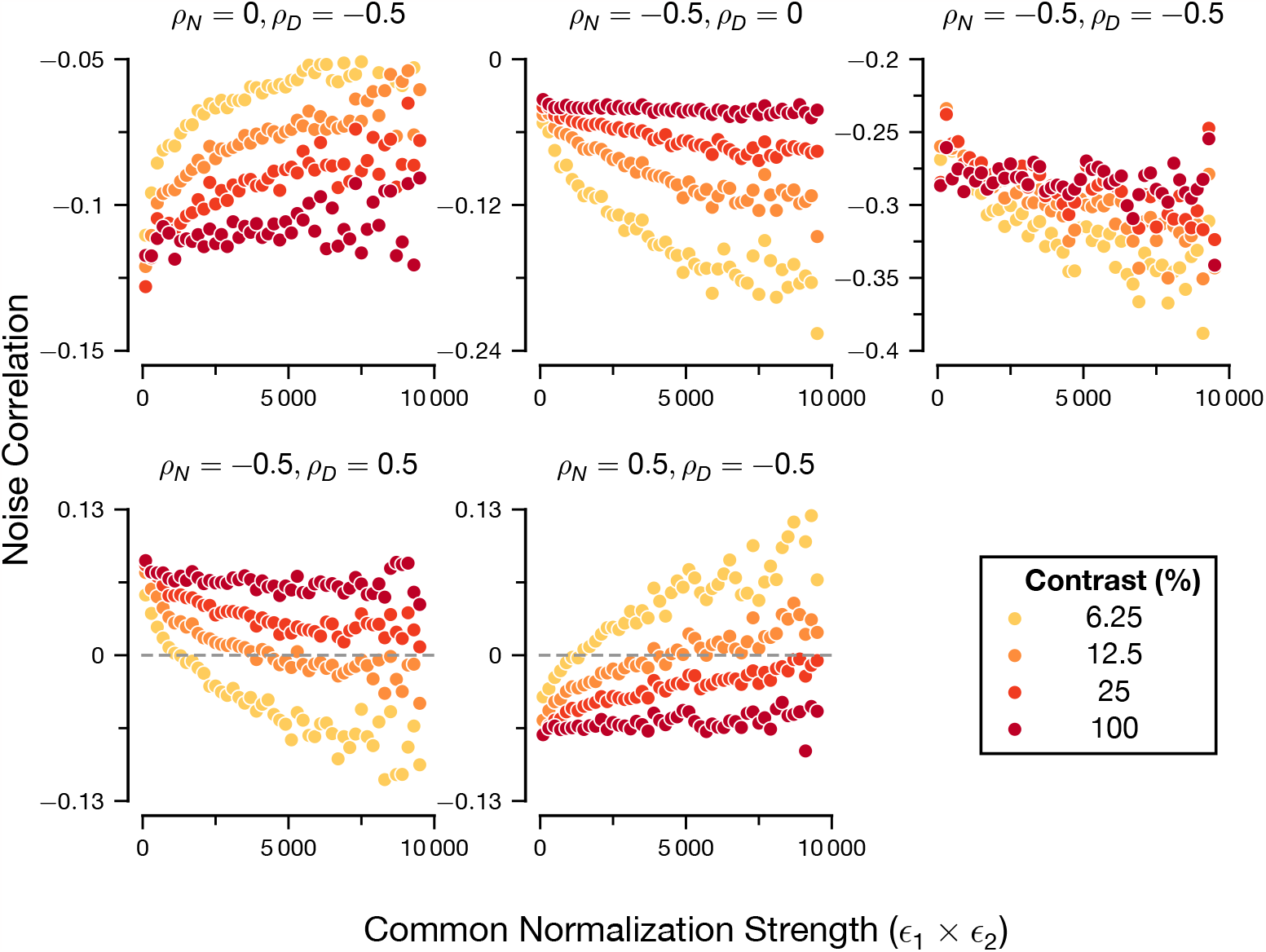
Relationship Between Noise Correlations and Denominator Strength. Expands upon Figure 2 to include cases where (*ρ*_*N*_, *ρ*_*D*_) can be negative or have opposite signs. Figure was created using the exact same method and synthetic dataset as Figure 2: see the caption in the main text for details.

**Figure S4:**
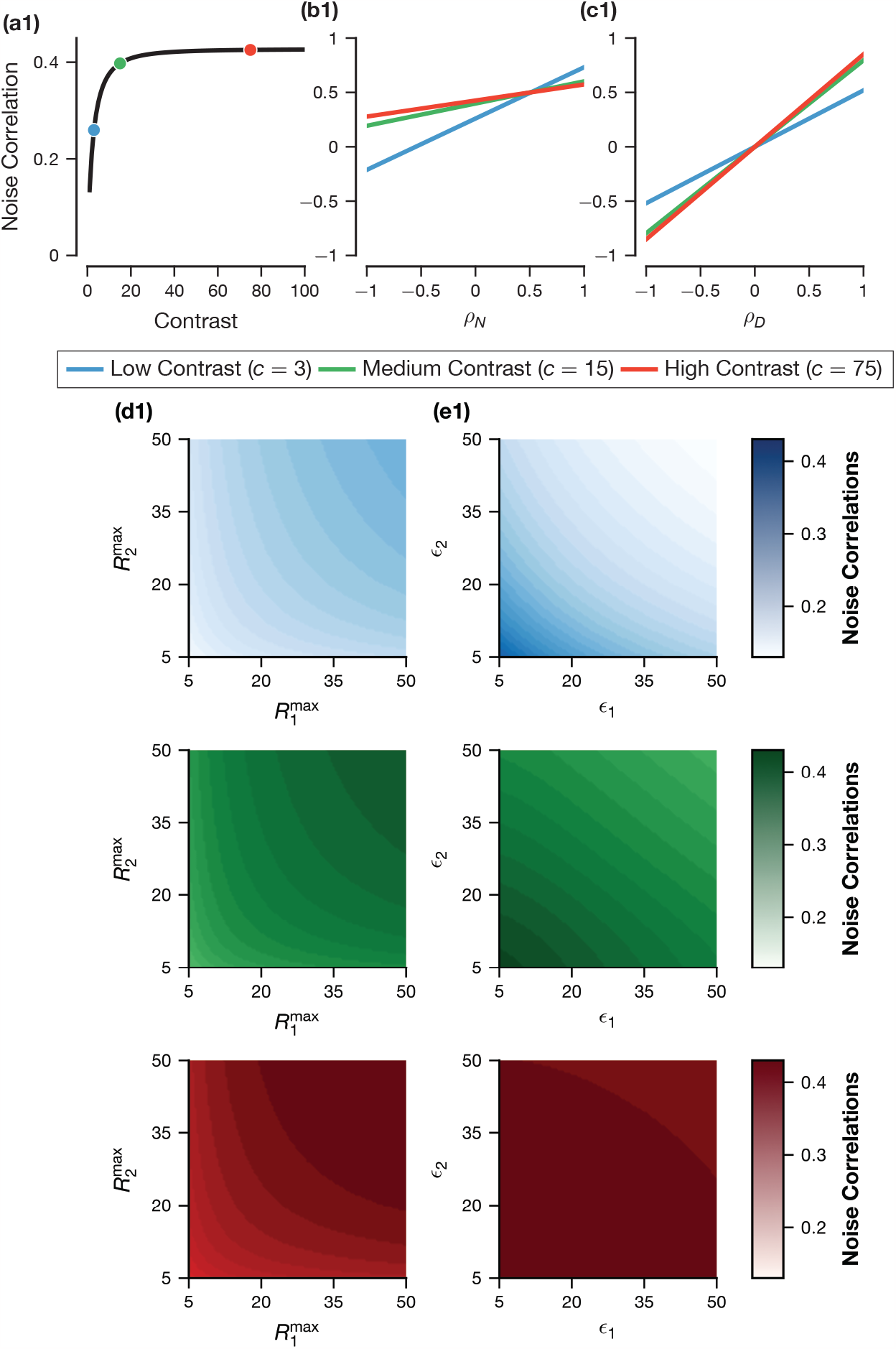
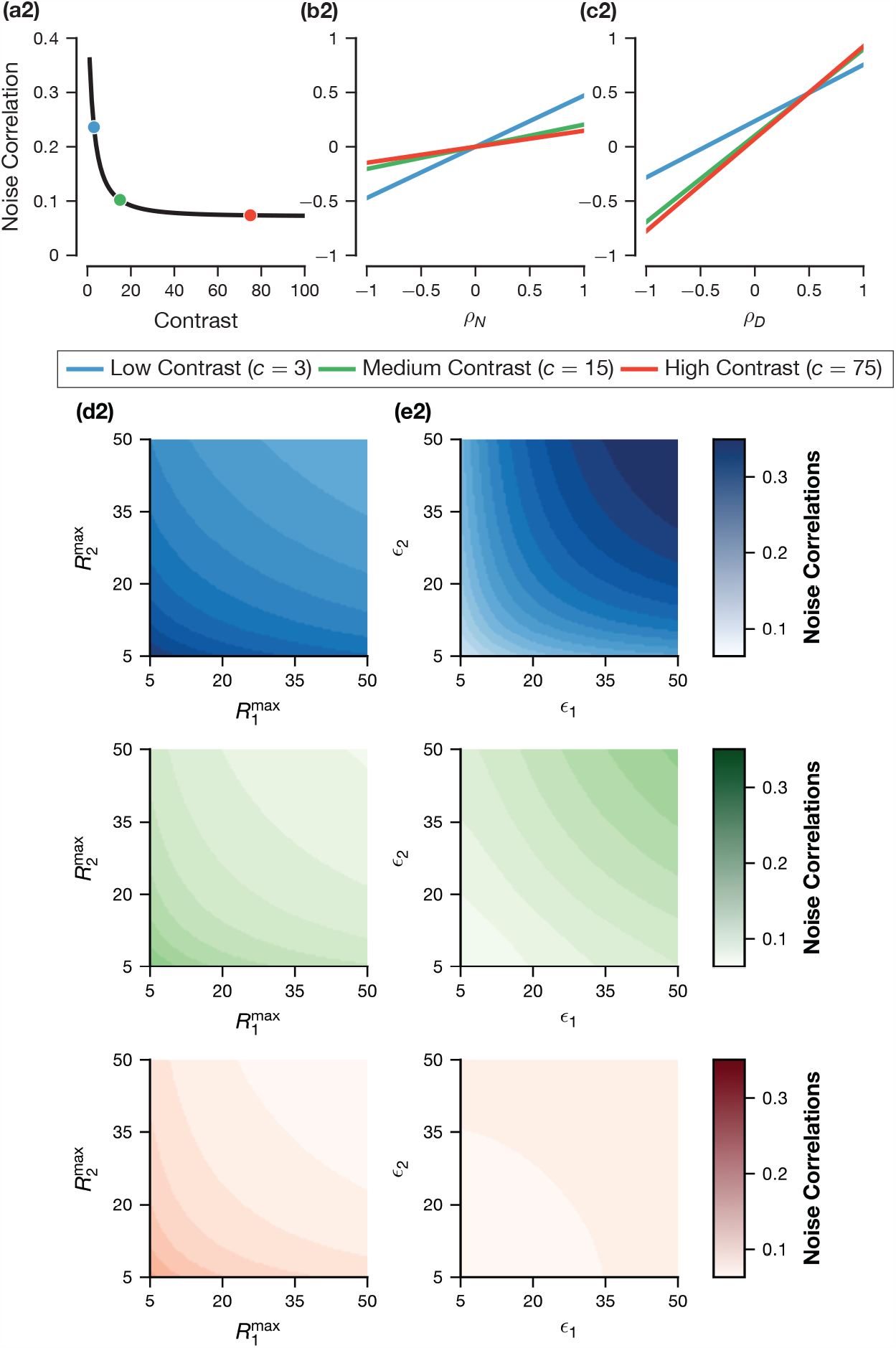
Relationship Between Noise Correlations and Ratio of Gaussians Parameters. Noise correlations in the model (Equation 7) can be modulated by stimulus strength (i.e., contrast), the correlation parameters of the model (*ρ*_*N*_, *ρ*_*D*_) and the parameters of the normalization model, in this case 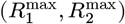 and (ϵ_1_, ϵ_2_) (see 2). To understand these effects in isolation, we looked at how Equation 7 changed with respect to each parameter, while keeping the other parameters constant. We illustrate with noise correlations that increase with contrast (a1-e1), and correlations that decrease with contrast (a2-e2). (a) Dependence of noise correlations on contrast. Three contrast levels that are fixed in the other panels are shown. (b) Dependence of noise correlations on *ρ*_*N*_.(c) Dependence of noise correlations on *ρ*_*D*_. (d) Dependence of noise correlations on 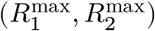, shown as a contour plot with the shade of color indicating noise correlation level. Different colors indicate different contrast levels as shown in the legend. (e) Dependence of noise correlations on (ϵ_1_, ϵ_2_).(a1-e1) uses the following parameters (when not fixed): 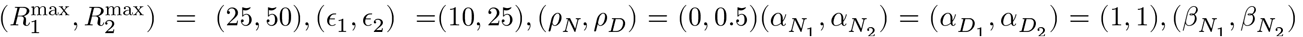 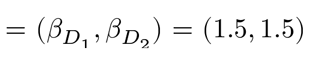(a2-e2) uses the same parameters except with (*ρ*_*N*_, *ρ*_*D*_) = (0.5, 0). Contrast levels were {1,…,100}.

